# Diversity and function of methyl-coenzyme M reductase-encoding archaea in Yellowstone hot springs revealed by metagenomics and mesocosm experiments

**DOI:** 10.1101/2022.08.18.504445

**Authors:** Mackenzie M. Lynes, Viola Krukenberg, Zackary J. Jay, Anthony J. Kohtz, Christine A. Gobrogge, Rachel L. Spietz, Roland Hatzenpichler

## Abstract

Metagenomic studies on geothermal environments have been central in recent discoveries on the diversity of archaeal methane and alkane metabolism. Here, we investigated the methanogenic populations inhabiting terrestrial geothermal features in Yellowstone National Park (YNP) by combining amplicon sequencing with metagenomics and mesocosm experiments. Detection of gene amplicons of methyl-coenzyme M reductase subunit A (*mcrA*) indicated a wide diversity of Mcr-encoding archaea across geothermal features with differing physicochemical regimes. From three selected hot springs we recovered twelve Mcr*-*encoding metagenome assembled genomes (MAGs) affiliated with lineages of cultured methanogens as well as *Candidatus* (*Ca*.) Methanomethylicia, *Ca*. Hadesarchaeia, and Archaeoglobi. These MAGs encoded the potential for hydrogenotrophic, aceticlastic, or hydrogen-dependent methylotrophic methanogenesis, or anaerobic short-chain alkane oxidation. While Mcr-encoding archaea represented a minor fraction of the microbial community of hot springs, mesocosm experiments with methanogenic precursors resulted in stimulation of methanogenic activity and the enrichment of lineages affiliated with *Methanosaeta* and *Methanothermobacter* as well as with uncultured Mcr-encoding archaea including *Ca*. Korarchaeia, *Ca*. Nezhaarchaeia, and Archaeoglobi. Altogether, we revealed that diverse Mcr-encoding populations with the metabolic potential to produce methane from different precursors persist in the geothermal environments of YNP. This study highlights the importance of combining environmental metagenomics with laboratory-based experiments to expand our understanding of uncultured Mcr-encoding archaea and their potential impact on microbial carbon transformations in geothermal environments and beyond.

## Introduction

Methane (CH_4_) is a climate active gas and an integral component in the global carbon cycle. The majority of biogenic methane is generated in anoxic environments by methanogenic archaea (1-3) that conserve energy by reducing low molecular weight substrates such as H_2_/CO_2_, acetate, or methylated compounds to CH_4_ (4-8). The final step in methanogenesis, the conversion of methyl-coenzyme M and coenzyme B into CH_4_, is catalyzed by the methyl-coenzyme M reductase (MCR) complex. This enzyme also catalyzes the reversible reaction, the activation of CH_4_ in anaerobic methane-oxidizing archaea (9) that, together with methanogens, control methane fluxes from anoxic environments, impacting global methane emissions to the atmosphere (1, 2). Currently, all cultured methanogens belong to lineages within the Euryarchaeota, and their physiology and biochemistry has been studied for decades (10-15). However, recent metagenomic studies discovered genes encoding the MCR complex in metagenome-assembled genomes (MAGs) from a variety of archaeal groups including *Candidatus* (*Ca*.) Methanofastidiosales, *Ca*. Nuwarchaeales, some members of the Archaeoglobi (16-18), as well as members of the TACK superphylum *Ca*. Methanomethylicia (*Ca*. Verstraetearchaeota), *Ca*. Korarchaeia, *Ca*. Bathyarchaeia, and *Ca*. Nezhaarchaeia (15, 19-22). Some of these *mcrA* genes have been shown to be transcribed in situ (23-26). Additionally, some MAGs contain highly divergent genes homologous to *mcr* (21, 22, 27-32). These *mcr*-like genes encode an alkyl-coenzyme M reductase (ACR) complex, which was experimentally shown to activate short-chain alkanes (*i*.*e*., ethane, propane, butane) in *Ca*. Synthrophoarchaeum, *Ca*. Argoarchaeum, and *Ca*. Ethanoperedens (27, 29, 31). However, as most newly discovered Mcr-encoding archaea are yet uncultured, their proposed methanogenesis and anaerobic methane/alkane metabolism awaits further experimental evaluation. MAGs representing these archaea have frequently been recovered from anoxic and often high temperature geothermal environments, such as deep-sea hydrothermal vents and terrestrial hot springs (18-21, 27, 30, 33). Methanogenesis in the geothermal system of Yellowstone National Park (YNP, Wyoming, USA), was initially studied in 1980 (34) and to date, three strains of the hydrogenotrophic methanogen *Methanothermobacter thermoautotrophicus* represent the only methanogens isolated from YNP (34, 35). However, *mcrA* and 16S rRNA genes affiliated with methanogenic archaea and *mcr*-containing MAGs have been repeatedly recovered from geothermal features across YNP (20, 21, 24, 33, 36), demonstrating the potential for a microbial methane cycle involving diverse archaeal lineages (SI Fig. 1).

In this study, we further explored the potential for methanogenesis in YNP by (1) combining *mcrA* gene amplicon sequencing with aqueous geochemical measurements to identify methanogenic populations across 100 previously uncharacterized geothermal features, and (2) employing metagenomics and methanogenic mesocosm experiments to investigate the metabolic potential and activity of methanogenic populations in three selected hot springs. We describe the taxonomic diversity of *mcrA* genes detected across contrasting physicochemical conditions, detail the metabolic potential of *mcr*-containing MAGs, and reveal the responses of Mcr-encoding archaea to methanogenic precursor substrates.

## Materials and Methods

### Selection of geothermal features and sample collection

A survey of *mcrA* gene diversity and aqueous geochemistry was conducted on 100 geothermal features including hot springs and mud pots with temperature between 18 - 94 °C and pH between 1.7 - 9.4 distributed across four geothermal regions in YNP (SI Fig. 1). Because many of these geothermal features are not included in the YNP Research Coordination Network (http://rcn.montana.edu/), unique sample identifiers are used in this study, which indicate the area, feature, and DNA sample (*e*.*g*., LCB003.1 denotes DNA sample 1 from feature 003 in the Lower Culex Basin). Sediments and microbial mats for *mcrA* gene amplicon sequencing and shotgun metagenomics were collected in 2017 and 2018 using a stainless-steel cup, homogenized, and frozen immediately until DNA extraction. A slurry of sediment and water (1:9) for mesocosm experiments was obtained in 2019 (SI Table 1, 2, 3) using a stainless-steel cup, transferred into a glass bottle, and homogenized. A 10 mL subsample was frozen immediately to preserve material for DNA extraction (environmental sample) before the bottle was sealed headspace-free. The slurry was transported in a heated container (∼50 °C), placed at in situ temperature within 4 h and used to set up mesocosm experiments within 12 h of retrieval.

**Table 1.** Statistics for twelve *mcr*-containing MAGs reconstructed from LCB metagenomes. Taxonomy for MAGs was assigned using 18 marker proteins (SI Table 4). Average GC content, completeness, coverage, and redundancy were determined with CheckM. Total abundance was inferred based on the average coverage and size of each MAG relative to all MAGs in the metagenome. * indicates *mcrA* was split across two scaffolds. Quality was assessed according to Bowers et al. 2017. Cov., coverage; Compl., completeness; Redund., redundancy; Abund., total abundance.

| MAG | Taxonomy | Size (Mbp) | Contigs | Genes | Cov. | GC (%) | Compl. (%) | Redund. (%) | Abund. (%) | tRNAs | rRNAs 16S,23S,5S | *mcrA* | Quality |
| --- | --- | --- | --- | --- | --- | --- | --- | --- | --- | --- | --- | --- | --- |
| LCB019-065 | *Methanolinea* | 1.78 | 27 | 1,847 | 587.6 | 57.9 | 98.7 | 0 | 2.33 | 17 | 1,1,1 | 1 | Medium |
| LCB019-055 | *Methanothermobacter* | 1.55 | 217 | 1,841 | 31.9 | 49.8 | 96.5 | 0.46 | 0.11 | 19 | 1,2,3 | 2 | High |
| LCB019-064 | *Methanotherix* | 1.66 | 72 | 1,674 | 491.4 | 54.6 | 100 | 0 | 1.82 | 16 | 0,0,2 | 1 | Medium |
| LCB019-061 | Methanomassiliicoccales | 1.41 | 237 | 1,501 | 24.3 | 44.3 | 87.61 | 3.23 | 0.08 | 13 | 1,0,1 | 1 | Medium |
| LCB019-004 | *Ca. Methanomethylicia* | 1.17 | 216 | 1,433 | 33.8 | 53.7 | 91 | 4.52 | 0.09 | 11 | 0,0,0 | 3 | Medium |
| LCB019-019 | *Ca. Methanosuratus* | 1.09 | 182 | 1,312 | 87.4 | 56.0 | 87.5 | 1.01 | 0.21 | 13 | 0,0,0 | 1 | Medium |
| LCB019-026 | *Ca. Methanomethylovorales* | 1.48 | 111 | 1,638 | 200.3 | 48.8 | 99.1 | 1.25 | 0.66 | 17 | 0,2,1 | 1* | Medium |
| LCB024-024 | *Ca. Methanomethylovorales* | 1.41 | 221 | 1,673 | 64.9 | 49.6 | 88.5 | 2.1 | 0.24 | 16 | 0,0,1 | 1* | Medium |
| LCB003-007 | *Ca. Methanomethyloarchaeales* | 1.03 | 67 | 1,243 | 26.7 | 27.9 | 93.5 | 0 | 0.11 | 15 | 0,0,1 | 1 | Medium |
| LCB024-038 | *Ca. Methanomethyloarchaeales* | 1.17 | 53 | 1,368 | 18.7 | 27.8 | 99.1 | 0.07 | 0.06 | 18 | 1,1,1 | 1 | High |
| LCB024-003 | Archaeoglobi | 1.16 | 179 | 1,399 | 87.2 | 45.7 | 88.5 | 1.31 | 0.27 | 18 | 0,1,0 | 2 | Medium |
| LCB024-034 | *Ca. Hadesarchaeia* | 0.73 | 188 | 945 | 62.6 | 66 | 72.1 | 0.93 | 0.12 | 15 | 0,0,0 | 1 | Medium |

### Physicochemical measurements, aqueous geochemistry, and elemental analysis

Temperature and pH were recorded in the water column of geothermal features using a thermocouple and portable pH meter. Water samples for aqueous geochemistry were collected and analyzed for dissolved iron, sulfide, and gases (O_2_, CH_4_, CO_2_) as previously described (37-39). Water samples for elemental analysis, anions, total carbon, inorganic carbon, non-purgeable organic carbon, and total nitrogen were processed by the Environmental Analytical Lab (Montana State University). Details in SI Materials and Methods.

### Mesocosm experiments

Mesocosm experiments with material from hot springs LCB003, LCB019, and LCB024 were prepared in an anoxic glove box (N_2_/CO_2_/H_2_; 90/5/5 %). Under constant stirring, 10 mL aliquots of sediment slurry were distributed into 25 mL serum bottles using serological plastic pipettes. Mesocosms were set up with the following treatments: acetate, formate, H_2_, H_2_ plus CO_2_ and bicarbonate, methanol, monomethylamine, methanol plus H_2_, monomethylamine plus H_2_, paraformaldehyde (killed control), bromoethanesulfonate (methanogenesis inhibitor), and no amendment. Two sets of triplicates per treatment were performed: (1) with bacterial antibiotics streptomycin (inhibitor of protein synthesis) and vancomycin (inhibitor of peptidoglycan synthesis) and (2) without antibiotics. Liquid amendments were added to a final concentration of 5 mM except for paraformaldehyde (5% v/v) and antibiotics (50 mg/L). Serum bottles were sealed with butyl rubber stoppers and aluminum crimps before the headspace was exchanged with N_2_ (99.999%) at 100 kPa for 5 min and set to 200 kPa N_2_. H_2_ and CO_2_ were added by exchanging an equal volume of N_2_ for a final concentration of 50% H_2_ and/or 20% CO_2_. Mesocosms were incubated at in situ temperatures: 74 °C (LCB003), 55 °C (LCB019), 72 °C (LCB024). Mesocosm headspace gas composition was monitored by subsampling 2 mL using a gastight syringe for analysis with a Varian gas chromatograph (GC; model CP2900) equipped with a dual-channel thermal conductivity detector system with Ar and He as carrier gases. Triplicate mesocosms were terminated simultaneously when the first replicate reached a CH_4_ plateau or exhausted supplied H_2_. At this time, a 0.5 mL slurry subsample was pelleted (12,000 g for 5 minutes) and frozen for DNA extraction. Mesocosms with low or no methane production including controls were terminated after 43 (LCB003), 35 (LCB019), or 55 (LCB024) days.

### DNA extraction and gene amplification

DNA was extracted from environmental samples (1 mL) and mesocosm samples (pellet from 0.5 mL) using the FastDNA Spin Kit for Soil (MP Biomedicals, Irvine, CA) following the manufacturer’s guidelines. *mcrA* genes were amplified with primer set mlas-mod-F/mcrA-rev-R (40, 41) from environmental DNA extracts. Archaeal and bacterial 16S rRNA genes were amplified with the updated Earth Microbiome Project primer set 515F and 806R (42-44) from DNA extracts of the mesocosm experiment. Amplicon libraries were prepared as previously described (45) and sequenced by Laragen Inc. (Culver City, CA) or the Molecular Research Core Facility at Idaho State University (Pocatello, ID) using an Illumina MiSeq platform with 2 × 300 bp (*mcrA* amplicon library) and 2 × 250 bp (16S rRNA amplicon library) paired end read chemistry. Details in SI Materials and Methods.

### Amplicon sequence analysis

Both 16S rRNA and *mcrA* gene reads were processed using QIIME 2 version 2020.2 (46). In short, primer sequences were removed from demultiplexed reads using cutadapt (47) with error rate 0.12, reads were truncated (145 bp forward, 145 bp reverse and 260 bp forward, 200 bp reverse for 16S rRNA and *mcrA* datasets, respectively), filtered, denoised and merged in DADA2 with default settings (48). 16S rRNA gene amplicon sequence variants (ASVs) were taxonomically classified with the sklearn method and the SILVA 132 database (49). *mcrA* gene ASVs were assigned a taxonomy using vsearch with a minimum identity of 70% and no consensus classification against a reference database of representative near-full length *mcrA* genes encompassing the diversity of publicly available *mcrA*. Contamination was removed using the R package decontam (50). The *mcrA* gene dataset was curated by removing ASVs ≤400 bp and non-*mcrA* gene ASVs as identified by evaluating the top hits of a blastx search against the NCBI NR database. Samples with less than 5,000 reads or 10,000 reads for the 16S rRNA and *mcrA* gene dataset, respectively, were excluded from further analyses. Diversity metrics and Bray-Curtis dissimilarity were calculated with the R packages phyloseq (51) and vegan (52).

### Metagenome sequencing, assembly, annotation

Metagenomes were generated at the Joint Genome Institute (JGI) from 10 ng (LCB003.1) and 100 ng (LCB019.1 and LCB024.1) DNA, and raw reads were processed according to JGI’s analysis workflow (see SI Materials and Methods). Quality controlled reads were assembled using SPAdes 3.11.1 (53) with options -m 2000, -k 33,55,77,99,127 -meta. Assembled scaffolds ≥2,000 bp were binned with six implementations of four different programs including, Maxbin v2.2.4 (54), Concoct v1.0.0 (55), Metabat v2.12.1 (56), and Autometa v1 (57). Bins generated from each program were refined with DAS_Tool (58) and bin quality statistics were determined with CheckM (59). MAGs were assigned alphanumerical identifiers (*e*.*g*., LCB003-007 indicates bin 7 from feature 003 in the Lower Culex Basin) and MAGs containing at least one *mcrA* gene (>300 nt) and a complete or near-complete set of *mcrABGCD* were considered in this study. For these Mcr-encoding MAGs, annotations provided by the IMG/M-ER pipeline v7 (60) for genes associated with methanogenesis pathways, coenzyme and cofactor biosynthesis, energy conservation, and beta-oxidation, were manually evaluated using analysis of gene neighborhoods, NCBI BLASTP, NCBI’s Conserved Domain Database, TMHMM, InterPro, and the hydrogenase classifier HydDB (61-64). See SI Data 3 for a complete list of gene annotations relevant to this study. Amino acid identity (AAI) values were computed with compareM using aai_wf and – proteins and taxonomies were assigned with GTDB-Tk v1.2.0 (65) and GTDB release 207 (66).

### Phylogenetic analyses

A set of 18 single-copy marker proteins (67, 68) detected in LCB Mcr-encoding MAGs and selected publicly available archaeal reference genomes (SI Table 4) were aligned using MUSCLE (69), trimmed with trimAL (70) with 50% gap threshold, and concatenated. A maximum likelihood phylogenetic tree was reconstructed with IQ-tree2 v2.0.6 (71) using the final concatenated alignment of 3,916 positions, LG+F+R10 model, and 1000 ultrafast bootstraps.

McrA from LCB metagenomes (>100 aa), abundant ASVs (140 aa), and publicly available references were aligned with MAFFT-linsi (72), trimmed with trimAL with 50% gap threshold and used for maximum likelihood phylogenetic analysis with IQtree2 with LG+C60+F+G model and 1000 ultrafast bootstraps.

### Data availability

16S rRNA gene and *mcrA* gene amplicon data as well as *mcr-*containing MAGs are deposited at NCBI under BioProject PRJNA859922 (SI Table 5). Metagenomes are available on IMG/M (JGI) under IMG Genome IDs 3300028675 (LCB003), 3300031463 (LCB019), and 3300029977 (LCB024).

## Results and Discussion

### Survey of *mcrA* genes across physicochemically contrasting geothermal features

The presence and diversity of Mcr-encoding populations was assessed in 100 geothermal features of YNP via *mcrA* gene amplicon sequencing. Gene amplicons were recovered from 66 sediment and/or microbial mat samples spanning 39 geothermal features located in the Lower Culex Basin (LCB; 61 samples, 35 features), the Mud Volcano Region (MVR; 4 samples, 3 features), and the White Creek Area (WCA; 1 sample, 1 feature). These features were characterized by a wide range of temperature (22 - 86.3 °C), pH (2.40 - 9.77), and dissolved methane (40 - 1,784 nM), oxygen (<13 - 771 µM), and sulfide (<2 - 27 µM) (Fig. 1; SI Fig. 1, SI Data 1).

**Figure 1.**
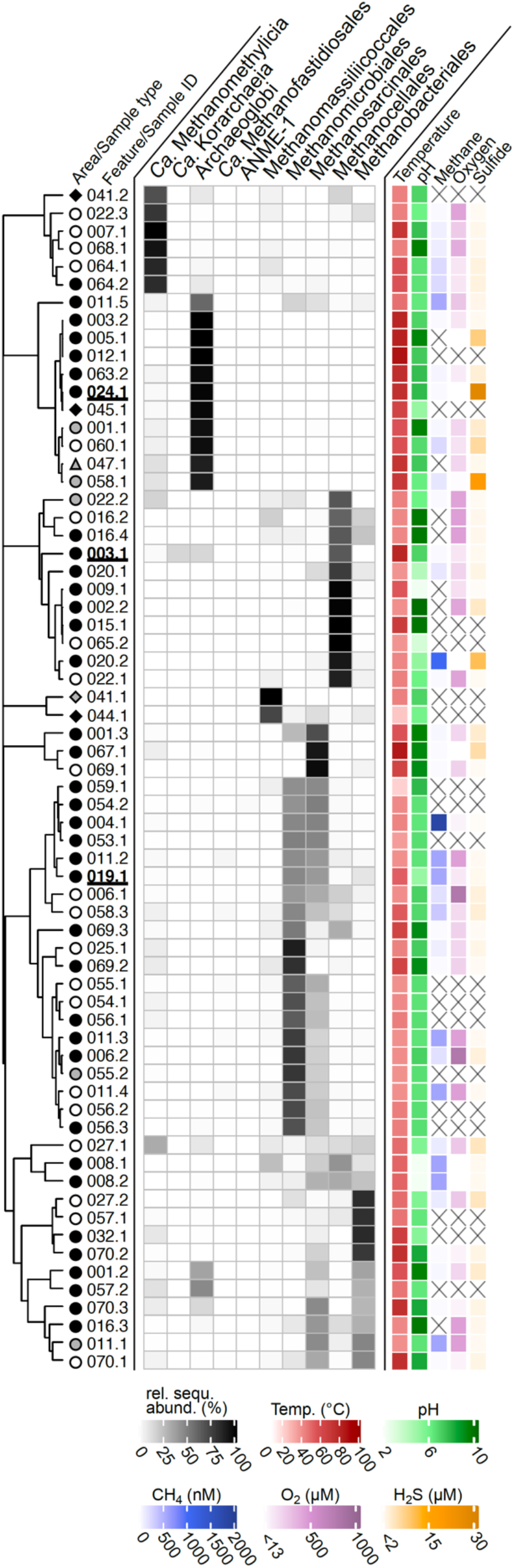
Diversity of *mcrA* genes detected in 66 samples from 39 geothermal features of YNP. Relative sequence abundance of *mcrA* gene amplicons affiliated with abundant lineages (relative sequence abundance >1% in at least one sample). Samples selected for metagenomics are underlined in bold. Samples were collected from geothermal features (identified by numbers) in the Lower Culex Basin (LCB, circle), Mud Volcano Region (MVR, diamond), and White Creek Area (WCA, triangle) and consisted of either sediment (black), microbial mat (white), or a mixture of sediment and mat material (grey). Physicochemical parameters of the geothermal water were recorded at the time of sample collection. X: no data available. Clustering based on Bray-Curtis dissimilarity using relative sequence abundance data of the presented lineages. No correlative trends between taxonomic affiliation of *mcrA* genes and physicochemistry were observed (SI Fig. 3). See SI Data 1 for details.

Generally, the *mcrA*-containing microbial community in each geothermal feature was composed of a small number (1-21) of *mcrA* ASVs with >1% relative sequence abundance. The alpha diversity of *mcrA* tended to decrease with increasing temperature (SI Fig. 2, SI Data 1), a trend consistent with previous results based on 16S rRNA gene diversity in geothermal environments (36, 73, 74). The *mcrA-*containing populations detected across samples included both confirmed methanogens (Methanomassiliicoccales, Methanosarcinales, Methanomicrobiales, Methanocellales, and Methanobacteriales) and lineages with proposed but untested methane/alkane metabolism (Archaeoglobi, *Ca*. Methanofastidiosales, *Ca*. Methanomethylicia, and *Ca*. Korarchaeia) (Fig. 1, SI Data 1). Confirmed methanogens dominated most samples (46/66) and were frequently identified in geothermal features with moderate temperatures (<60 °C). Other Mcr-encoding lineages prevailed at elevated temperatures (>60 °C) and were exclusively detected in LCB024.1, LCB058.1, and LCB063.2 (SI Fig. 2, 3, SI Data 1). Particularly, Archaeoglobi-affiliated *mcrA* genes dominated at high temperatures (≥70 °C) and elevated concentrations of dissolved sulfide (≥3 µM (SI Fig. 2)), which are conditions similar to those environments in which Archaeoglobi were previously detected (18, 21, 23, 75). Notably, *Ca*. Korarchaeia were present at high relative sequence abundance (16%) in LCB003.1. Anaerobic methane-oxidizing archaea (ANME-1) as well as *Ca*. Methanofastidiosales were detected only with low relative sequence abundance (<2 and <4%, respectively) (SI Data 1).

Overall, this survey of *mcrA* genes indicated that taxonomically diverse *mcrA-*containing archaea exist across a wide range of physicochemical regimes in the geothermal environments of YNP, particularly in the LCB geothermal area. As primer-based diversity surveys are inherently biased, we note that the primer set used in this study has historically been widely applied to amplify *mcrA* genes of Euryarchaeota origin (41). While these primers bind to currently known *mcrA* genes of *Ca*. Methanomethylicia and *Ca*. Korarchaeia without mismatches, multiple mismatches to *mcrA* genes of other lineages exist (*e*.*g*., *Ca*. Nezhaarchaeia). Consequently, our amplicon-based gene survey likely underrepresented certain *mcrA* genes and underestimated *mcrA* gene diversity. To further investigate the methanogenic communities in the LCB, metagenomics and mesocosm experiments were conducted with material from hot springs LCB003, LCB019, and LCB024 characterized by elevated temperatures (47 - 73 °C) and circumneutral pH (3.0 - 7.8) (SI Table 1, 2, 3, SI Fig. 4).

### *mcrA* gene diversity and *mcr*-containing MAGs recovered from the LCB

In the hot springs selected as main study sites, abundant *mcrA* ASVs were related to confirmed methanogens in LCB019 and LCB003, and archaea with proposed methane/alkane metabolism in LCB024 and LCB003 (SI Table 6, SI Data 1). Environmental metagenomics recovered ten medium and two high quality MAGs (76) encoding McrA and a complete/near-complete MCR complex, which ranged in size from 0.73 - 1.78 Mbp and estimated completeness from 72 - 100% (Table 1). According to phylogenomic analysis of 18 archaeal single copy genes (SI Table 4), four MAGs belonged to lineages of previously cultured methanogens: *Methanothermobacter* (LCB019-055), *Methanomassiliicoccales* (LCB019-061), *Methanothrix* (LCB019-064), and *Methanolinea* (LCB019-065) while eight MAGs belonged to lineages of proposed methanogens or methane/alkane-oxidizing archaea: *Ca*. Methanomethylicia (LCB019-004, -019, -026, LCB024-024, -038, LCB003-007), Archaeoglobi (LCB024-003), and *Ca*. Hadesarchaeia (LCB024-034) (Fig. 2, SI Fig. 5). Interestingly, *Ca*. Methanomethylicia related MAGs were recovered from all three hot spring metagenomes indicating that members of this lineage exist across various physicochemical regimes. In contrast, MAGs affiliated with lineages of confirmed methanogens were only identified in LCB019, as initially reflected by *mcrA* amplicons. In total, 12 near-complete (≥500 aa) and 24 partial (100 - 499 aa) McrA sequences were recovered from the metagenome assemblies, suggesting that the Mcr-encoding MAGs reconstructed here do not reflect the full diversity and metabolic potential of the Mcr-encoding populations present. According to phylogenetic analysis, McrA proteins were categorized into MCR-type and ACR- type (21, 77) and affiliated with McrA of confirmed methanogens (group I), proposed methanogens (group II), or with McrA-like proteins of proposed alkane-metabolizing archaea (group III) (22, 27) (Fig. 2). Overall, the *mcrA* genes and Mcr-encoding MAGs recovered via metagenome sequencing confirmed the diversity of *mcrA-*containing archaea detected via amplicon sequencing and extended it by detecting Methanomassiliicoccales, *Ca*. Nezhaarchaeia, and *Ca*. Hadesarchaeia (Fig. 2).

**Figure 2.**
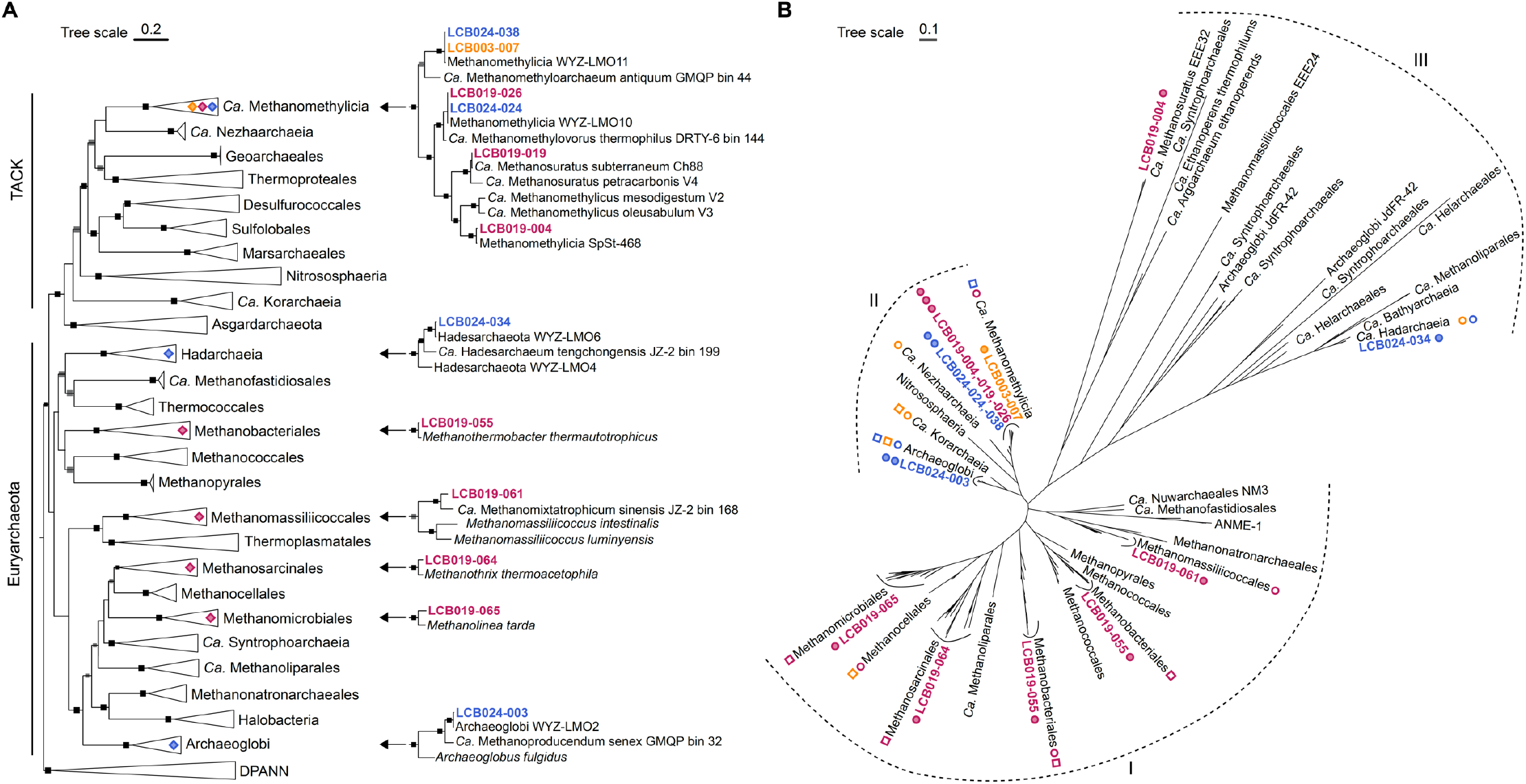
Phylogenetic tree of Mcr-encoding MAGs and McrA. (A) Maximum-likelihood tree, inferred with IQtree and the best-fit LG+F+R10 model, using a concatenated set of 18 conserved arCOGs (SI Table 4). Squares indicate ultrafast bootstrap values of 100 (black) and 95-99 (gray). Diamonds indicate lineages with Mcr-encoding MAGs detected in this study and shown in detail. (B) Maximum-likelihood tree, inferred with IQtree and the LG+C60+F+G model, from the amino acid alignment of McrA. Filled circles: McrA identified in a MAG, open circles: metagenomic McrA (unbinned, >100 aa), open squares: abundant *mcrA* ASVs (>1% relative sequence abundance). For details see SI Table 6. Dashed line indicates previously proposed McrA/AcrA groups (Wang et al., 2019, 2021): I) McrA from methanogens and ANME (MCR-type), II) McrA from TACK lineages (MCR-type), III) McrA-like from proposed and experimentally confirmed alkane oxidizing archaea (ACR-type). Colors: orange, LCB003, magenta, LCB019, blue, LCB024.

### Potential for methane and alkane metabolism in *mcr*-containing MAGs

Four MAGs were affiliated with lineages of confirmed hydrogenotrophic, aceticlastic, and hydrogen-dependent methylotrophic methanogens and encoded *mcrA* genes related to those of cultured methanogens (group I). LCB019-065 and LCB019-055 shared amino acid identity (AAI) values of 80% and 98% with cultured representatives of the hydrogenotrophic methanogens *Methanolinea* and *Methanothermobacter*, respectively. Congruently, both MAGs encoded the genes required for generating methane from H_2_ and CO_2_, including the complete Wood-Ljungdahl Pathway (WLP), methyl-H_4_M(S)PT:coenzyme M methyltransferase (Mtr) complex, F_420_-reducing hydrogenase (Frh), methyl-viologen-reducing hydrogenase (Mvh) (incomplete in LCB019-065), and energy-converting hydrogenase (Ehb, LCB019-055; Ech, LCB019-065) (Fig. 3, SI Discussion). Additionally, a complete formate dehydrogenase complex (FdhABC) was encoded in LCB019-65 and while LCB019-055 encoded FdhAB, FdhC was not detected. Consistently, cultured representatives of *Methanolinea* utilize formate as a substrate for methanogenesis while those of *Methanothermobacter* do not (78, 79). LCB019-064 showed AAI values of 90% to the aceticlastic methanogen *Methanothrix thermoacetophila* and encoded all genes necessary for aceticlastic methanogenesis including the Mtr complex and acetyl-CoA decarbonylase/synthase:CO dehydrogenase complex (ACS/CODH) (Fig. 3, SI Discussion). LCB019-061 shared AAI of 59% with cultured *Methanomassiliicoccus* sp., suggesting it may represent a novel lineage within the Methanomassiliicoccales. Consistent with a hydrogen-dependent methylotrophic methanogenesis lifestyle of Methanomassiliicoccales isolates, LCB019-061 encodes methyltransferases (SI Fig. 6) but lacks the WLP and a complete Mtr complex. A *mtrH* gene encoded in proximity to methyltransferase corrinoid activation protein (*ramA*) suggests LCB019-061 may reduce unknown methylated substrates to methane (19, 75).

**Figure 3.**
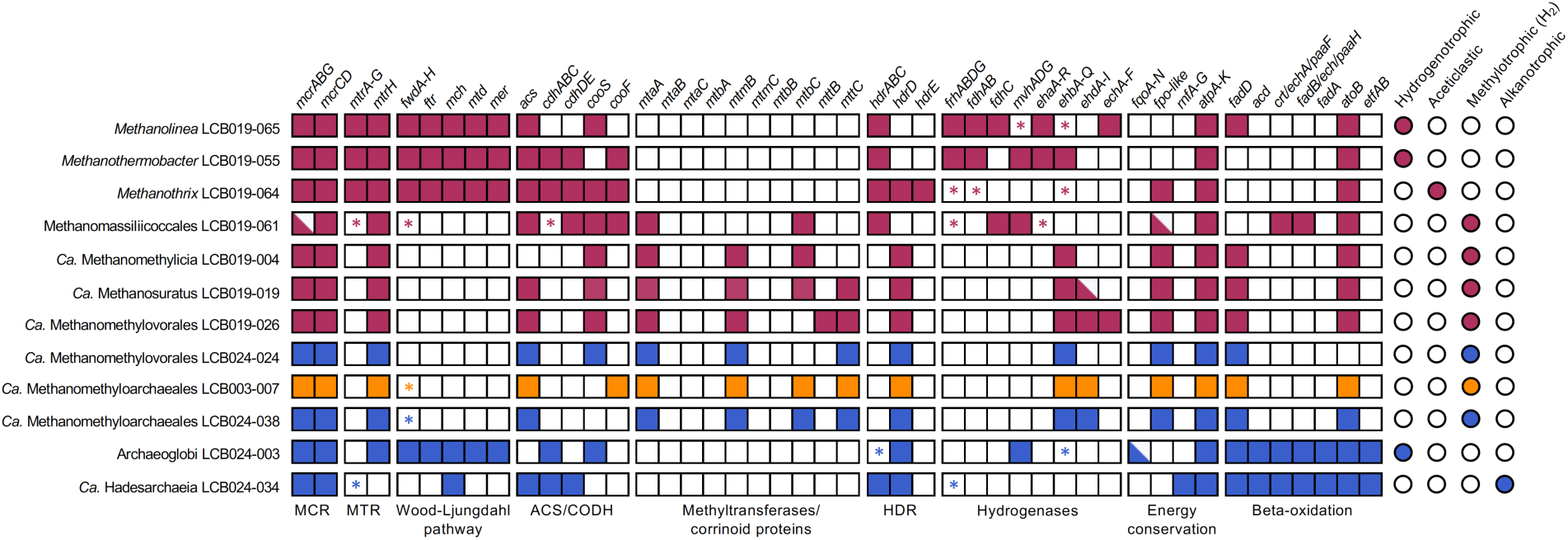
Methanogenic potential of the twelve Mcr-encoding MAGs. Squares indicate gene/gene set detected (filled), gene/gene set not detected (open) or gene set partially detected with the majority of genes present (half filled). * indicates only one gene in a gene set detected. Circles indicate the methane/alkane metabolism predicted for each MAG based on the gene repertoire. Colors: orange, LCB003, magenta, LCB019, blue, LCB024. A complete list of genes described in this figure and their abbreviations is reported in SI Data 3.

Six MAGs shared high AAI values (>96%) with MAGs of *Ca*. Methanomethylicia and encoded an McrA affiliated with those of other *Ca*. Methanomethylicia MAGs (group II). Consistent with *Ca*. Methanomethylicia MAGs proposed to perform hydrogen-dependent methylotrophic methanogenesis (19), the six MAGs lack the WLP and a complete Mtr complex but encode a variety of methyltransferases including methanol:coenzyme M methyltransferase (*mtaA)*, monomethylamine methyltransferase (*mtmB*), and dimethylamine corrinoid (*mtbC*) and/or trimethylamine corrinoid protein (*mttC*). LCB019-026 additionally encoded a trimethylamine methyltransferase (*mttB*). A methyltransferase subunit H of the Mtr complex, *mtrH*, encoded near other corrinoid protein and methyltransferase genes (SI Data 3) suggests that methane may be formed from unknown methylated substrates (19, 75). Although methylamine-specific cobamide:coenzyme M methyltransferase (*mtbA*) was absent, MtaA could substitute for the activity of MtbA (SI Fig. 6) (80). Thus, all six *Ca*. Methanomethylicia MAGs contain the gene repertoire needed for hydrogen-dependent methylotrophic methanogenesis (Fig. 3, SI Fig. 6, SI Discussion). In addition, LCB019-004 encoded a second McrA, that clustered with the McrA-like proteins of ethane-oxidizing archaea *Ca*. Ethanoperedens and *Ca*. Argoarchaeum (McrA group III, ACR/ECR type) and a recently recovered MAG of *Ca*. Methanosuratus (25, 77) proposed to perform ethanogenesis or ethane oxidation via an unknown pathway (77). This indicates that anaerobic methane/alkane metabolism within the *Ca*. Methanomethylicia may be more diverse than previously anticipated.

The Archaeoglobi affiliated MAG LCB024-003 showed low AAI values (65%) to *Archaeoglobales* isolates, which are all non-methanogenic sulfate-reducers. Instead, LCB024-003 shared high AAI values (>98%) to Mcr-encoding Archaeoglobi MAGs of proposed hydrogenotrophic methanogens (WYZ-LMO10, SJ34) or hydrogen-dependent methylotrophic methanogens (*Ca*. M. hydrogenotrophicum) (21, 33, 75). Consistently, its two partial McrA (192 and 193 aa) cluster with McrA of other proposed methanogenic Archaeoglobi (group II) (Fig. 2) (18, 21). LCB024-003 encodes genes required for hydrogenotrophic methanogenesis including the WLP pathway, hydrogenase Mvh, and a F_420_H_2_:quinone oxidoreductase complex (*fqo*DHIF) which may substitute for Frh to generate reduced F_420_ as previously suggested (23, 81, 82); however, a complete Mtr complex was not detected. In contrast to *Ca*. M. hydrogenotrophicum, LCB024-003 encodes a truncated 5,10-methylenetetrahydromethanopterin reductase (*mer*) while *mtaABC* were not identified, suggesting it is unable to use methanol for methanogenesis (23). Although LCB024-003 encodes the beta-oxidation pathway, other genes typically associated with short-chain alkane oxidation including an ACR-type MCR, ACS/CODH complex, and methyltransferases were absent. Hence, unlike the MAGs of *Ca*. Polytropus marinifundus and JdFR-42 (18, 21), LCB024-003 may not represent an anaerobic alkane oxidizer (83) and instead may utilize the beta-oxidation pathway for long chain fatty acid metabolism as has been shown for *Archaeoglobus fulgidus* (84). Further, genes encoding dissimilatory sulfate reduction (*sat, aprAB, dsrABC*) present in some *mcr-*containing Archaeoglobi MAGs (*Ca*. M. dualitatem (23)) were not detected. Together, the genomic information from LCB024-003 suggests that this Archaeoglobi representative may live as a hydrogenotrophic methanogen (Fig. 3, SI Discussion).

LCB024-034 shared AAI values of >79% with other Mcr-encoding Hadesarchaeia (21, 75) and encoded a partial McrA (216 aa) related to the ACR-type proteins of Hadesarchaeia (group III) (27). Congruently with the hypothesis of short-chain alkane metabolism in *Ca*. Hadesarchaeia, LCB024-034 encoded the beta-oxidation pathway and an ACS/CODH complex. However, most genes encoding the WLP required for oxidizing activated alkanes to CO_2_ were missing (27). Thus, short-chain alkane metabolism in LCB024-034 remains speculative, awaiting further genomic and experimental data.

Together, the twelve *mcr-*containing MAGs reconstructed here reflect the potential for archaeal short-chain alkane-oxidation as well as hydrogenotrophic, aceticlastic, and hydrogendependent methylotrophic methanogenesis in geothermal environments of YNP. Further, these MAGs extend the genomic data available for future analysis of diversity and evolution of Mcr-encoding archaea and suggest geothermal environments are a promising source for the recovery of these archaea.

### Methanogenic activity and enrichment of methanogens in mesocosms

Mesocosm experiments were performed to reveal activity and enrichment of methanogens. Methane accumulation was monitored in the headspace of mesocosms under (1) close to in situ conditions (*i*.*e*., no amendment), (2) conditions favoring methanogenesis (*i*.*e*., substrate amendment), and (3) conditions inhibiting bacterial metabolism (*i*.*e*., antibiotics treatment) (SI Fig. 7). Mesocosms were also analyzed for enrichment in potential methanogenic populations via 16S rRNA gene amplicon sequencing. Abundant 16S rRNA gene ASVs (>1% relative sequence abundance) related to Mcr-encoding archaea amounted for ∼1% in LCB019 and <1% in LCB024 and LCB003, indicating that methanogens represent a minor fraction of the in situ community (SI Fig. 8, 9). However, in mesocosms from all three hot springs, methane production was observed under close to in situ conditions with strongly varying maximum methane yields (17,000, 1,900 and 150 ppm for LCB019, LCB024 and LCB003, respectively) (Fig. 4, SI Fig. 7). Substrate amendment had considerably different effects on methane production and, except for LCB019, mesocosm triplicates showed strong variation and long response times (20-40 days) likely due to an uneven distribution of initially low abundant methanogen cells across replicates. For LCB024, substrate amendment (particularly H_2_), appeared to suppress methane production, which may indicate that either hydrogenotrophic methanogens were not present, not active, or were outcompeted by other community members considering the shift in the microbial community (see SI Fig. 9). Antibiotic amendments resulted, on average across treatments, in increased methane production in mesocosms from LCB024 and LCB003, and a strong decrease in methane production in mesocosms from LCB019, indicative of substrate competition or metabolic interdependencies between methanogens and bacteria, respectively.

**Figure 4.**
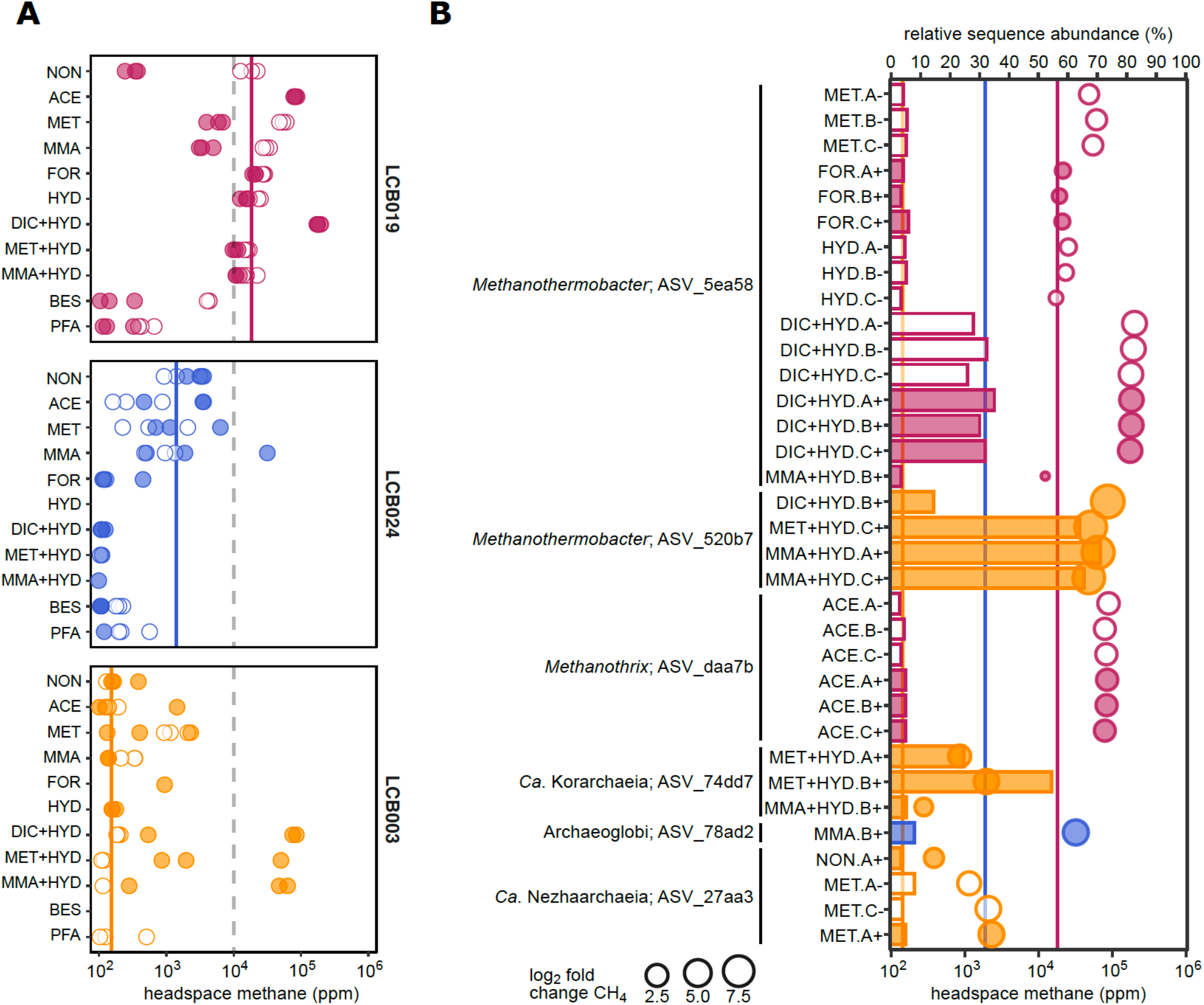
Methane production and enrichment of Mcr-encoding archaea in mesocosms. (A) Maximum methane produced in the headspace of mesocosms. Replicates measuring <100 ppm not shown. (B) Enrichment of 16S rRNA gene ASVs (>3% relative sequence abundance) affiliated with Mcr-encoding archaea across treatments (bars) paired with respective headspace methane yields (circles). Circle size proportional to the log2 fold change in methane yield between treatment and control (*i*.*e*., mesocosm under close to in situ condition) for each site. Dashed lines indicate 1% methane. Solid lines indicate average methane concentration in mesocosms under close to in situ conditions (no substrate, no antibiotics) for each site. Open symbols: without antibiotics; filled symbols: with antibiotics. Colors: orange, LCB003, magenta, LCB019, blue, LCB024. Abbreviations: NON, no amendment control; ACE, acetate; MET, methanol; MMA, monomethylamine; FOR, formate; HYD, hydrogen (H_2_); DIC, dissolved inorganic carbon (HCO_3_^-^+CO_2_); BES, bromoethanesulfonate (methanogenesis inhibitor); PFA, paraformaldehyde (killed control). Replicates indicated as A-C and +, with antibiotics or -, without antibiotics. Methane curves and extended relative abundance data for all mesocosm replicates are reported in SI Fig. 7-9 and SI Data 4.

To characterize the effect of substrate amendment on methanogenic populations, we analyzed ASVs related to Mcr-encoding archaea with enrichment >3% relative sequence abundance across treatments. H_2_ plus DIC (HCO_3_^-^+CO_2_) amended mesocosms from LCB019 showed rapid methane production, with highest maximum methane concentrations (>170,000 ppm) reached within 6 days (SI Fig. 7). These mesocosms were enriched (26-35%) in ASV_5ea58, identical to the 16S rRNA gene of MAG LCB019-055 as well as *Methanothermobacter thermautotrophicus*, a thermophilic hydrogenotrophic methanogen isolated from YNP (34, 35). Similarly, for LCB003, H_2_ plus DIC or methylated compounds resulted in the strongest stimulation of methanogenesis and most pronounced enrichment (up to 71%) of an ASV affiliated with *Methanothermobacter crinale* (ASV_520b7, 99.6% sequence identity). Notably, H_2_ amendment without DIC supply did not result in a comparable response, suggesting that in closed mesocosm systems hydrogenotrophic methanogens were limited by inorganic carbon, which unlikely occurs in situ where concentrations of aqueous CO_2_ were elevated (SI Table 2). For LCB019, acetate amendment resulted in elevated methane production and concomitant enrichment (3-5%) of ASV_daa7b, which shared high sequence similarity with the aceticlastic methanogen *Methanothrix thermoacetophila* (98%) and a 16S rRNA gene recovered from the LCB019 metagenome (100%; SI Data 4). MAG LCB019-064, related to *Methanothrix thermoacetophila* (89% AAI similarity) encoded the potential for methanogenesis from acetate and may represent the enriched *Methanothrix* sp. population. Thus, our mesocosm experiments complemented findings from metagenomics, confirming the potential for hydrogenotrophic methanogenesis by *Methanothermobacter* sp. and aceticlastic methanogenesis by *Methanothrix* sp. in LCB019 and revealing the potential for hydrogenotrophic methanogenesis by *Methanothermobacter* sp. in LCB003 (SI Table 6).

In addition to previously cultured methanogens, uncultured Mcr-encoding lineages were enriched. An ASV identified as *Ca*. Methanodesulfokores washburnensis (ASV_74dd7, 100% sequence identity) was highly abundant (25-54%) in two mesocosms from LCB003 amended with methanol, hydrogen, and antibiotics. A MAG of this *Ca*. Korarchaeia representative previously recovered from YNP encodes versatile metabolic capabilities including hydrogen-dependent methylotrophic methanogenesis from methanol (20). Methane yields in these mesocosms, while comparably low after 43 days (<2,000 ppm), were strongly elevated compared to the no amendment control of LCB003 (log2 fold change (FC) 3-4). *mcrA* and 16S rRNA genes of *Ca*. Methanodesulfokores washburnensis were also detected via amplicon and metagenome sequencing, confirming the presence of this lineage in LCB003 (Fig. 1, 2, SI Data 4). In one mesocosm from LCB024 amended with monomethylamine, stimulation of methanogenesis (log2 FC 4, 310,000 ppm) and enrichment (8%) of Archaeoglobi-affiliated ASV_78ad2 was observed.

The Archaeoglobi MAG LCB024-003 recovered from LCB024 encoded the potential for hydrogenotrophic methanogenesis while genes required for methylotrophic methanogenesis were not detected (Fig. 3). However, potential for methylotrophic methanogenesis has been described for some Archaeoglobi MAGs and the recovery of several Archaeoglobi related *mcrA* and 16S rRNA genes from LCB024 suggests that diverse Archaeoglobi populations are present, possibly including methylotrophic methanogens. An enrichment of a *Ca*. Nezhaarchaeia related ASV (ASV_27aa3) was highest (8%) in methanol amended mesocosms from LCB003 and cooccurred with elevated methane yields (log2 FC 3.5, 85,000 ppm), confirming the persistence of a *Ca*. Nezhaarchaeia population as previously detected by metagenomic 16S rRNA and *mcrA* genes (SI Data 2). Previously described *mcr*-containing *Ca*. Nezhaarchaeia MAGs encode the potential for hydrogenotrophic methanogenesis, and while no enrichment was detected in hydrogen amended mesocosms, microbially produced hydrogen may have facilitated limited methanogenic activity and enrichment of hydrogenotrophic methanogens in other mesocosms. *Ca*. Methanomethylicia related ASVs were detected in multiple mesocosms, however their enrichment remained low (<3%) (SI Data 4).

Overall, minor methanogenic populations, not or hardly detectable in hot springs via 16S rRNA gene or metagenome sequencing, were enriched in mesocosm experiments under selective methanogenic conditions. Specifically, methyl compounds favored enrichment of *Ca*. Korarchaeia, *Ca*. Nezhaarchaeia, or Archaeoglobi. However, further research is needed to decipher the metabolism of the enriched populations, their proposed methanogenic capacities, and potential metabolic interdependencies with other community members.

### Implications for methane cycling in YNP

This study explored the potential for methanogenesis in previously uncharacterized geothermal environments of YNP, primarily the LCB, and warrants further research into the magnitude of biological methane production in this area. We detected *mcrA* genes across a wide range of physicochemical regimes indicating the presence of diverse populations of Mcr-encoding archaea, including both confirmed methanogens and lineages proposed to engage in anaerobic methane/alkane cycling in geothermal environments. The methanogenic pathways encoded across *mcr*-containing MAGs suggests methanogenesis in LCB hot springs could proceed from different precursors including H_2_/CO_2_, acetate, and methyl compounds plus hydrogen. The genetic potential for hydrogen-dependent methylotrophic methanogenesis was encoded by the majority of MAGs, including *Ca*. Methanomethylicia and Methanomassiliicoccales, and was detected in all three hot springs, possibly reflecting prevalence of this metabolism in geothermal environments as previously proposed (75). While methanogenic populations accounted for a minor fraction of the microbial community, methanogenesis may proceed in situ as it was observed in mesocosms under close to in situ conditions. The potential for hydrogenotrophic and aceticlastic methanogenesis revealed by metagenomics was confirmed by the enrichment of *Methanothermobacter* and *Methanothrix* in mesocosms under selective substrate amendment. In situ, methanogenesis in hot springs is likely constrained by physicochemical regimes, substrate availability, and metabolic interdependencies. Methanogenic precursors may be supplied as metabolic intermediates of syntrophic communities (e.g., H_2_, acetate), products of respiration (e.g., CO_2_), or through geothermal alteration from the subsurface (e.g., H_2_, CO_2_) (85). As hot springs often present dynamic systems, methanogens may frequently respond with activity and growth to favorable conditions. This may be exemplified by *Methanothermobacter*’s capacity to rapidly respond, resulting in high activity and fast growth upon supply of H_2_/CO_2_, which it may sporadically or consistently encounter in situ (Fig. 4, SI Table 2, SI Fig. 7).

Although methanogens have previously been detected in different areas of YNP (34, 35), their environmental impact is not well understood. Methane is an important component of the gas flux in YNP (85-87) and the isotopic composition of gas emitted from geothermal features across YNP has suggested methane is primarily generated through abiogenic and/or thermogenic processes, while methanogenesis is not a significant source of methane (86). We detected varying concentrations of aqueous methane in geothermal features, but the source and fate of this methane is currently unknown. In general, methane emissions from terrestrial geothermal environments are not considered in estimates of the global atmospheric methane budget and little is known about their contribution to the global methane flux (1, 3, 14). YNP contains more than 14,000 geothermal features, the largest concentration in the world, making it a superior candidate for studying CH_4_ flux in these environments (88-90).

Environmental *mcrA* gene surveys and metagenomics aid in identifying areas of YNP where methanogenesis may occur. Subsequent quantification of in situ metabolic activities, including methane production rates, as well as deciphering the interplay between methanogens and methanotrophs will lead to a better understanding of the impact methanogens have on the local carbon cycle and their contribution to methane emissions from YNP’s geothermal environments.

## Conclusion

Uncultured Mcr-encoding lineages are globally distributed across a wide range of ecosystems and could play important roles in the biogeochemical carbon cycle (14, 18-24, 28, 30, 33). In this study, we described a previously unrecognized diversity of Mcr-encoding archaea in geothermal environments of YNP. Environmental metagenomics provided insights into the metabolic potential of these Mcr-encoding archaea and mesocosm experiments were essential for revealing their methanogenic activity and enrichment. The ability to enrich uncultured Mcr-encoding archaea presents an opportunity to recover their genomes and a step towards their cultivation. Future work, including experiments under close to in situ conditions and culture-dependent physiology and biochemistry studies, is essential for advancing our understanding of these enigmatic archaea.

## Acknowledgements

This study was funded through an NSF RII Track-2 FEC award DBI-1736255 (to R.H.). A portion of this research was performed under a Facilities Integrating Collaborations for User Science (FICUS) program award (proposal: 10.46936/fics.proj.2017.49972/6000002 to R.H.), and used resources at the Joint Genome Institute (JGI; https://ror.org/04xm1d337), which is a DOE Office of Science User Facility. JGI is sponsored by the Office of Biological and Environmental Research and operated under Contract No. DE-AC02-05CH11231. M.M.L. was supported in part by the Thermal Biology Institute and Montana State University’s Vice President’s Office of Research, Economic Development, and Graduate Education. We appreciate the U.S. National Park Service, in particular, Annie Carlson at the Yellowstone Center for Resources, for permitting work in Yellowstone National Park under permit number YELL-SCI-8010. We thank William Inskeep, Timothy McDermott, and Luke McKay for helpful discussions that informed field sampling efforts.

## Author contributions

M.M.L., V.K., and R.H. developed the study. M.M.L. and R.H. collected initial survey samples. M.M.L. and C.A.G. conducted geochemical analyses. Z.J.J. processed metagenomes, recovered MAGs, and provided bioinformatics support. A.J.K. and Z.J.J. performed phylogenetic analysis of MAGs. V.K. and Z.J.J. performed McrA phylogenetic analysis. M.M.L. and A.J.K reconstructed metabolic potential of MAGs. V.K. and M.M.L. prepared amplicon libraries and processed amplicon data. M.M.L., V.K., A.J.K, and R.L.S. designed and conducted mesocosm experiments. M.M.L. and V.K. devised the figures and wrote the initial manuscript with input from R.H. The final version was edited and approved by all authors.

## Supplementary Information

### Table of contents

**SI Data 1**. Environmental survey of physicochemistry and *mcrA* genes. See separate excel file.

**SI Data 2**. Additional information for Mcr-encoding MAGs. See separate excel file.

**SI Data 3**. Gene inventory for Mcr-encoding MAGs. See separate excel file.

**SI Data 4**. Mesocosm 16S rRNA gene amplicon data. See separate excel file.

**SI Table 1.**
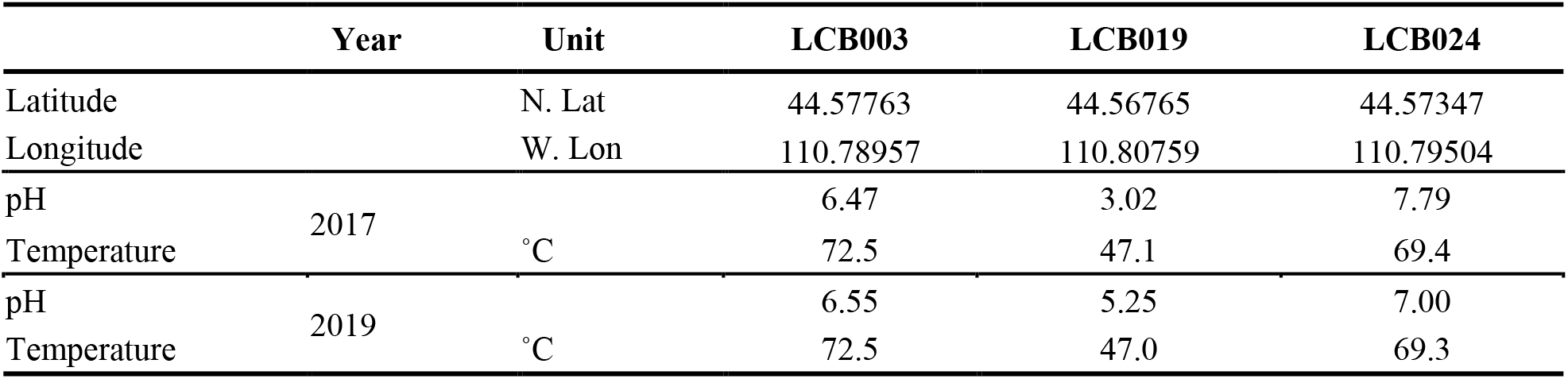
Location, pH, and temperature of main study sites. Data recorded at the time of sampling for metagenomics (2017) and mesocosm experiments (2019).

|  | Year | Unit | LCB003 | LCB019 | LCB024 |
| --- | --- | --- | --- | --- | --- |
| Latitude |  | N. Lat | 44.57763 | 44.56765 | 44.57347 |
| Longitude |  | W. Lon | 110.78957 | 110.80759 | 110.79504 |
| pH | 2017 |  | 6.47 | 3.02 | 7.79 |
| Temperature |  | °C | 72.5 | 47.1 | 69.4 |
| pH | 2019 |  | 6.55 | 5.25 | 7.00 |
| Temperature |  | °C | 72.5 | 47.0 | 69.3 |

**SI Table 2.** Aquatic geochemistry of main study sites (2017-2020). Data shown as ranges when applicable.

|  | Year | Unit | LCB003 | LCB019 | LCB024 |
| --- | --- | --- | --- | --- | --- |
| pH | 2017-2020 |  | 6.14 - 6.84 | 3.02 - 5.51 | 7.00 - 7.90 |
| Temperature |  | °C | 70.1 - 79.0 | 44.4 - 52.3 | 61.4 - 73.7 |
| CH<sub>4</sub> (aq) | 2019-2020 | nM | 40 - 211 | 470 - 576 | 30 - 77 |
| O<sub>2</sub> (aq) |  | μM | <13 - 117 | <13 - 171 | <13 - 131 |
| H<sub>2</sub>S (aq) |  | μM | 0.91 - 3.28 | 1.04 - 14.59 | 11.98 - 34.50 |
| Fe (II) |  | μM | 0 - 12 | 82 - 104 | 0 |
| total Fe (II, III) |  | μM | 0 - 20 | 97 - 99 | 0 - 1.1 |
| CO<sub>2</sub> (aq) |  | μM | 27 - 340 | 487 - 580 | 16 - 32 |
| PO<sub>4</sub><sup>3-</sup> | 2017 | μM | 0.98 | 6.11 | 0.79 |
| total N |  | μM | 19.70 | 56.78 | 9.55 |
| NH<sub>4</sub><sup>+</sup> |  | μM | 6.26 | 29.14 | 1.54 |
| NO<sub>2</sub><sup>-</sup> |  | μM | 0.05 | 0.04 | 0.24 |
| SO<sub>4</sub><sup>2-</sup> |  | mM | 0.14 | 0.36 | 0.15 |
| Cl<sup>-</sup> |  | mM | 8.03 | 4.88 | 8.53 |
| F<sup>-</sup> |  | mM | 1.28 | 0.95 | 1.53 |
| total C |  | mM | 2.57 | 2.56 | 2.96 |
| inorganic C |  | mM | 1.87 | 0.89 | 2.97 |
| non-purgeable organic C |  | mM | 0.18 | 0.30 | 0.07 |

**SI Table 3.** Elemental analysis of main study sites (2017). BD, below detection.

|  | Unit | LCB003 | LCB019 | LCB024 |
| --- | --- | --- | --- | --- |
| Ag | μM | 0.01 | BD | 0.06 |
| Al | μM | 8.34 | 44.48 | 9.45 |
| As | μM | 11.41 | 5.81 | 15.42 |
| Ba | μM | BD | 0.26 | BD |
| Be | μM | 1.00 | BD | BD |
| Bi | μM | 0.07 | BD | BD |
| Ca | μM | 52.40 | 67.37 | 28.07 |
| Cd | μM | BD | BD | BD |
| Co | μM | BD | BD | BD |
| Cr | μM | BD | BD | BD |
| Cu | μM | BD | BD | BD |
| Fe | μM | 1.37 | 6.98 | BD |
| Mg | μM | 11.73 | 22.83 | BD |
| Mn | μM | BD | 0.38 | BD |
| Mo | μM | 0.27 | BD | 1.72 |
| Ni | μM | BD | BD | BD |
| Pb | μM | 0.06 | BD | BD |
| Sb | μM | BD | BD | 0.17 |
| Se | μM | 0.28 | 0.09 | 0.11 |
| Sr | μM | BD | BD | BD |
| Ti | μM | 0.47 | BD | BD |
| Tl | μM | 0.01 | BD | BD |
| V | μM | 0.53 | 0.06 | 0.27 |
| Zn | μM | 0.53 | 0.16 | 0.11 |
| B | mM | 0.33 | 0.17 | 0.34 |
| K | mM | 0.13 | 0.35 | 0.25 |
| Li | mM | 0.33 | 0.23 | 0.40 |
| Na | mM | 10.25 | 5.09 | 12.16 |
| S | mM | 0.31 | 0.73 | 0.26 |
| Si | mM | 2.47 | 3.07 | 3.09 |

**SI Table 4.** NCBI BioSamples for MAGs under BioProject PRJNA859922.

| **MAG** | **Taxonomy** | **NCBI Taxonomy/Organism** | **BioSample** | **Accession** |
| --- | --- | --- | --- | --- |
| LCB019-065 | *Methanolinea* | *Methanolinea* sp. | SAMN29855412 | JANIHI000000000 |
| LCB019-055 | *Methanothermobacter* | *Methanothermobacter* sp. | SAMN29855422 | JANIHH000000000 |
| LCB019-064 | *Methanotherix* | *Methanotherix* sp. | SAMN29855467 | JANIHG000000000 |
| LCB019-061 | Methanomassiliicoccales | Methanomassiliicoccales archaeon | SAMN29855483 | JANHAK000000000 |
| LCB019-004 | *Ca. Methanomethylicia* | *Ca. Methanomethylicia* archaeon | SAMN29877625 | JANHAI000000000 |
| LCB019-019 | *Ca. Methanosuratus* | *Ca. Methanosuratincola* sp. | SAMN29877776 | JANIHJ000000000 |
| LCB019-026 | *Ca. Methanomethylovorales* | *Ca. Methanomethylicia* archaeon | SAMN29877790 | JANHAJ000000000 |
| LCB024-024 | *Ca. Methanomethylovorales* | *Ca. Methanomethylicia* archaeon | SAMN29851525 | JANHAL000000000 |
| LCB003-007 | *Ca. Methanomethyloarchaeales* | *Ca. Methanomethylicia* archaeon | SAMN29878106 | JANHAH000000000 |
| LCB024-038 | *Ca. Methanomethyloarchaeales* | *Ca. Methanomethylicia* archaeon | SAMN29851763 | JANHAM000000000 |
| LCB024-003 | Archaeoglobi | Archaeoglobi archaeon | SAMN29809817 | JANFND000000000 |
| LCB024-034 | *Ca. Hadesarchaeia* | *Ca. Hadesarchaea* archaeon | SAMN29854158 | JANIHF000000000 |

**SI Table 5.**
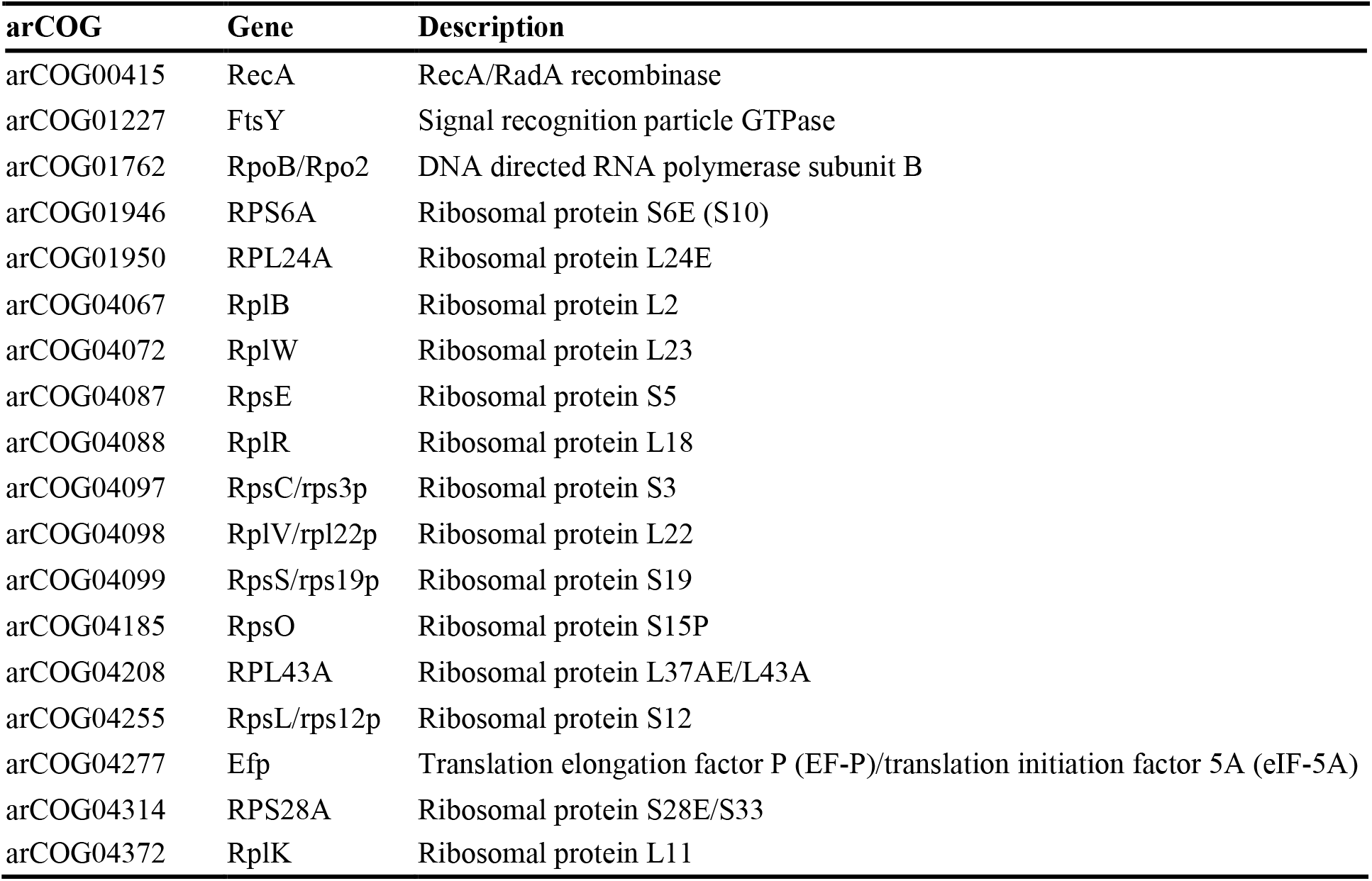
Marker genes used in phylogenomic analysis of MAGs.

**SI Table 6.**
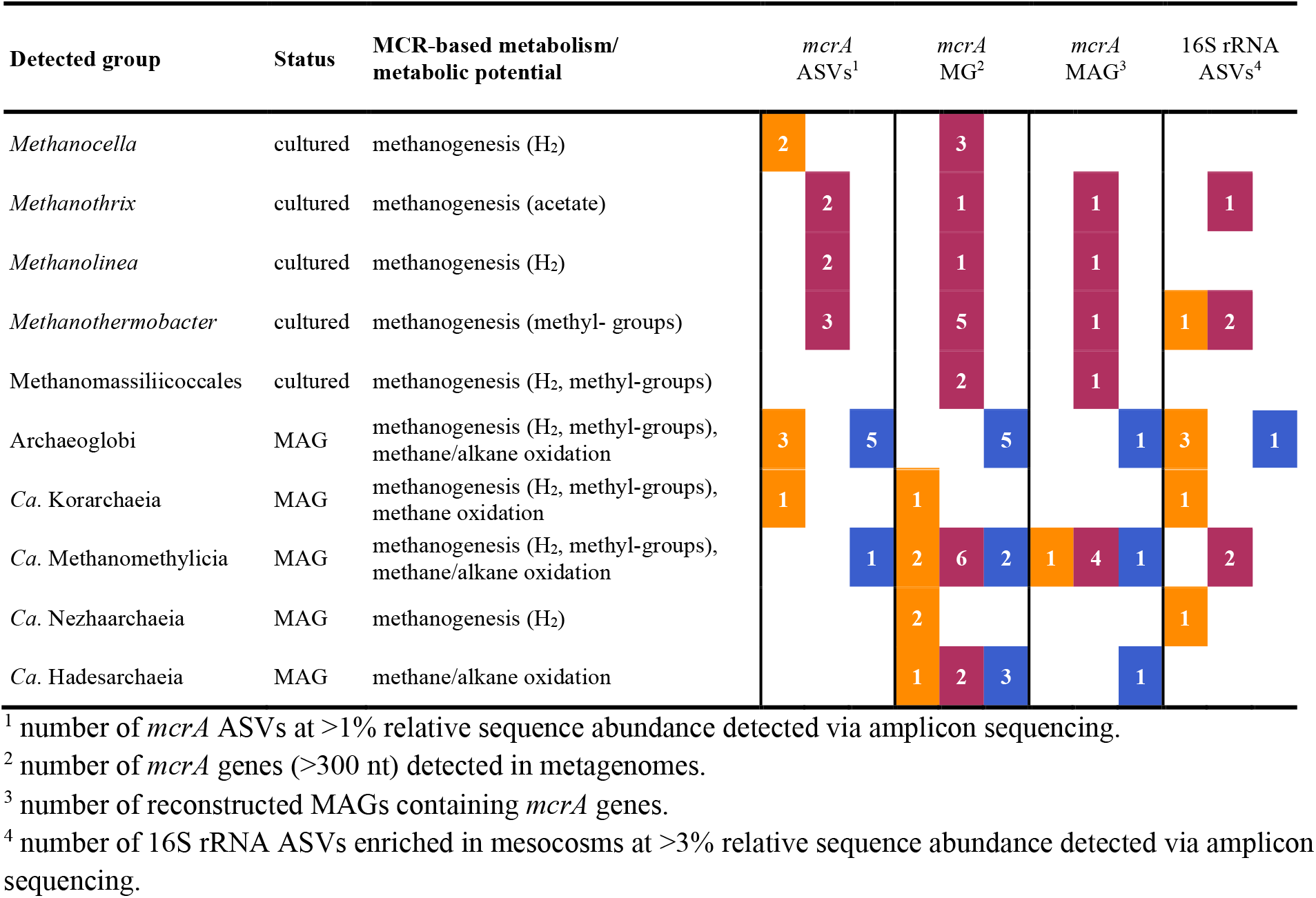
Overview of Mcr-encoding lineages detected in this study. Color coding by site: orange, LCB003, magenta, LCB019, blue, LCB024.

**SI Figure 1.**
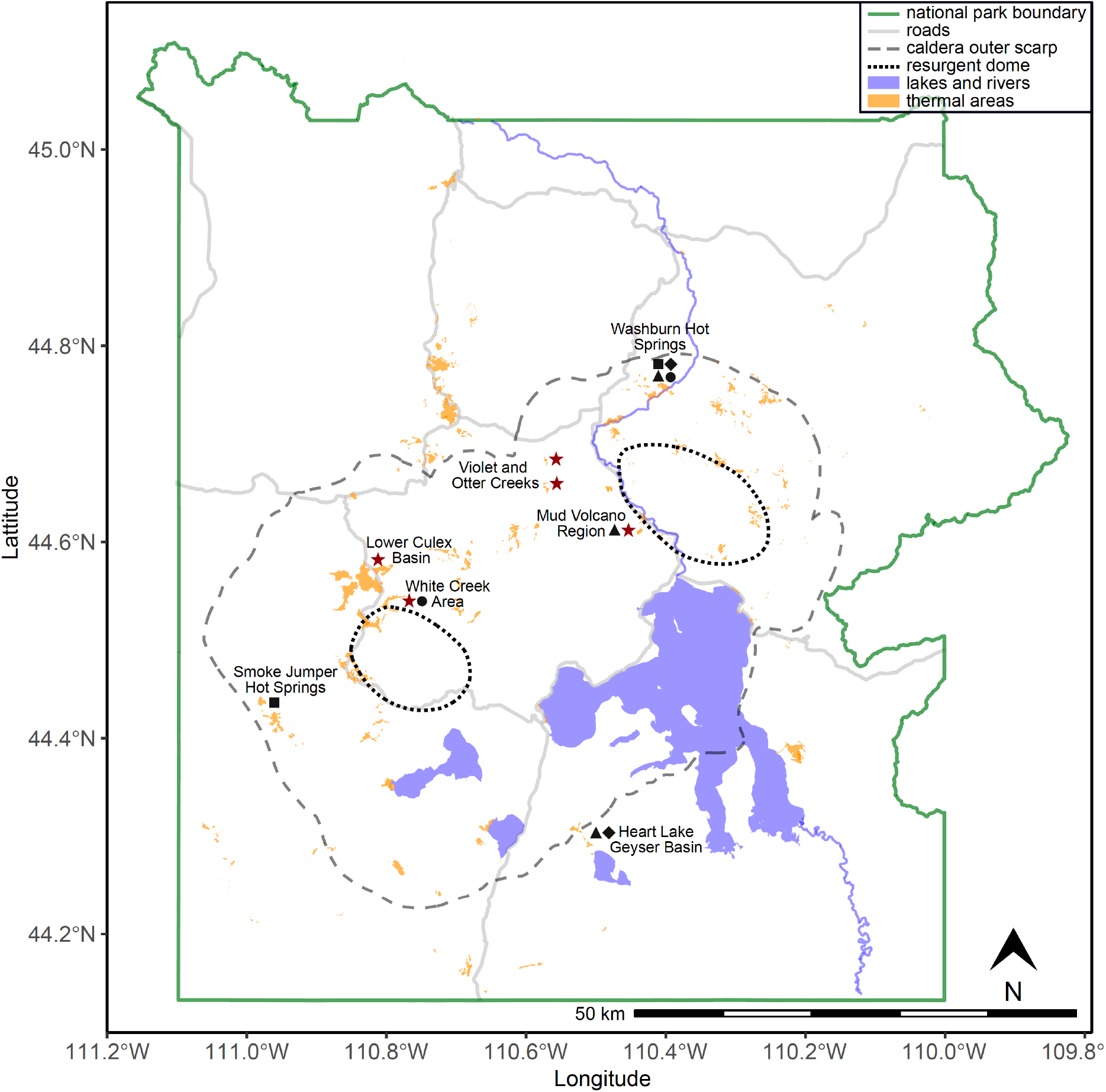
Yellowstone National Park map highlighting areas with potential for methanogenesis. Stars mark areas investigated in this study. Symbols indicate where *mcrA* genes (triangle), 16S rRNA genes of methanogens (diamond), Mcr-encoding MAGs (square), or methanogen isolates (circle) have been recovered. Map was generated in R with data from the Wyoming State Geological Survey (https://www.wsgs.wyo.gov/pubs-maps/gis).

**SI Figure 2.**
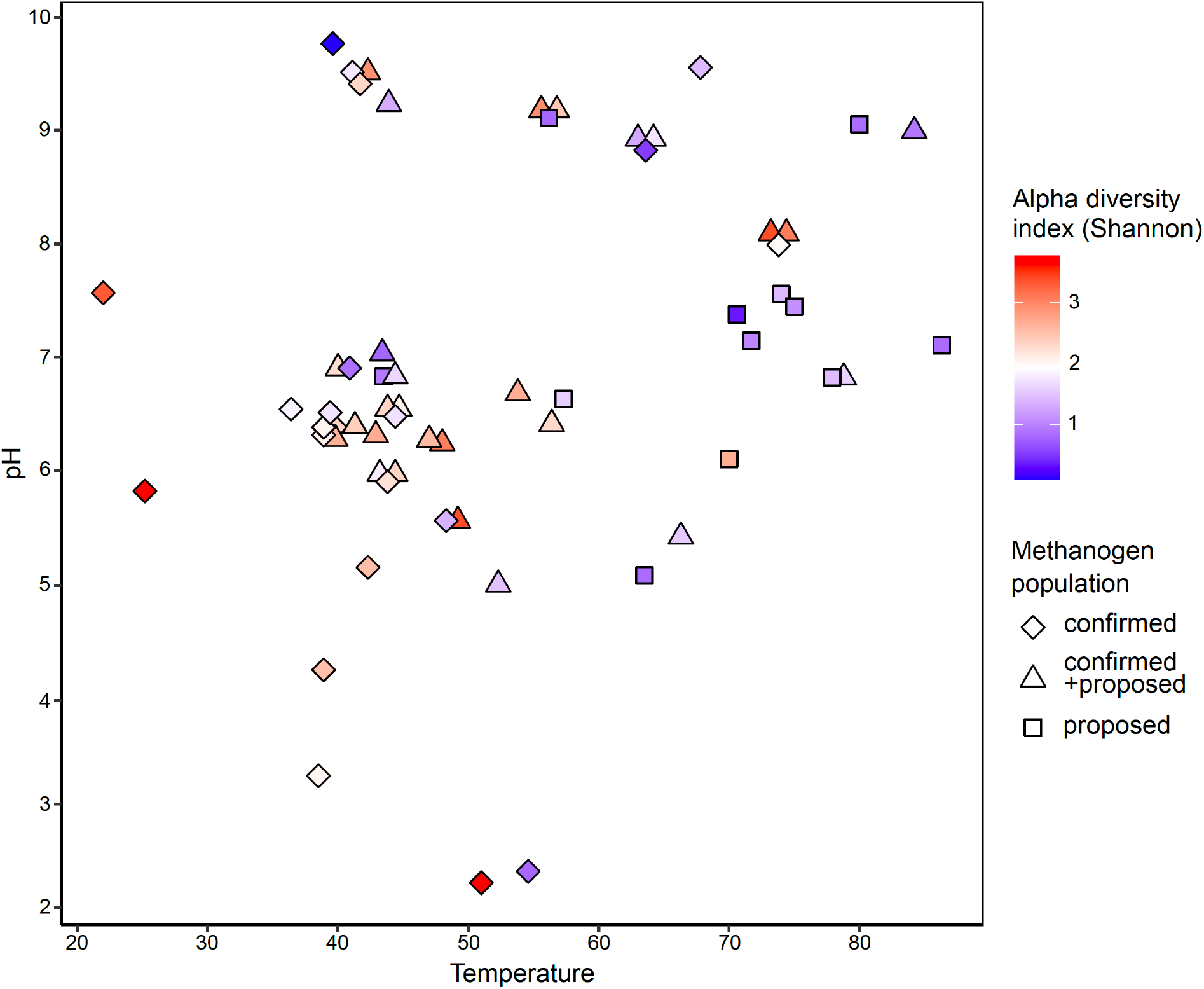
Alpha diversity of *mcrA* gene ASVs across temperature and pH. Most samples were retrieved from geothermal features with circumneutral pH and moderate to elevated temperatures (35-80°C). *mcrA* alpha diversity (Shannon index) indicates a trend towards decreased richness at elevated temperatures. Mcr-encoding communities tend to be primarily composed of confirmed methanogens at moderate temperatures and proposed methanogens or methane/alkane oxidizing archaea at elevated temperatures.

**SI Figure 3.**
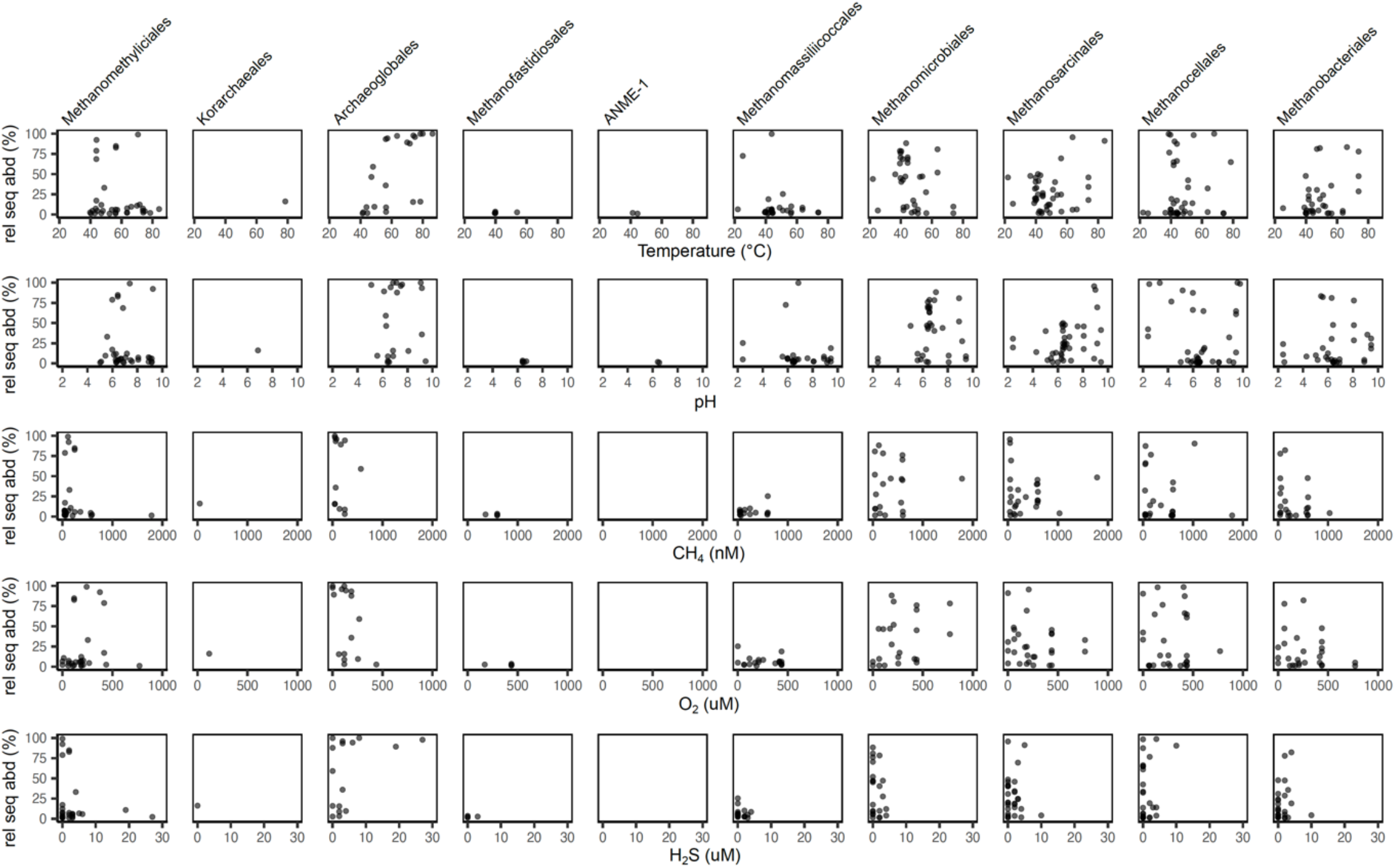
Distribution of Mcr-encoding archaea across physicochemical regimes. Mcr-encoding lineages detected via *mcrA* gene amplicon sequencing with relative abundances >1%. Aqueous CH_4_, O_2_, and H_2_S data not available for all samples. Detection limit for O_2_ and H_2_S was 13 µM and 2 µM, respectively.

**SI Figure 4.**
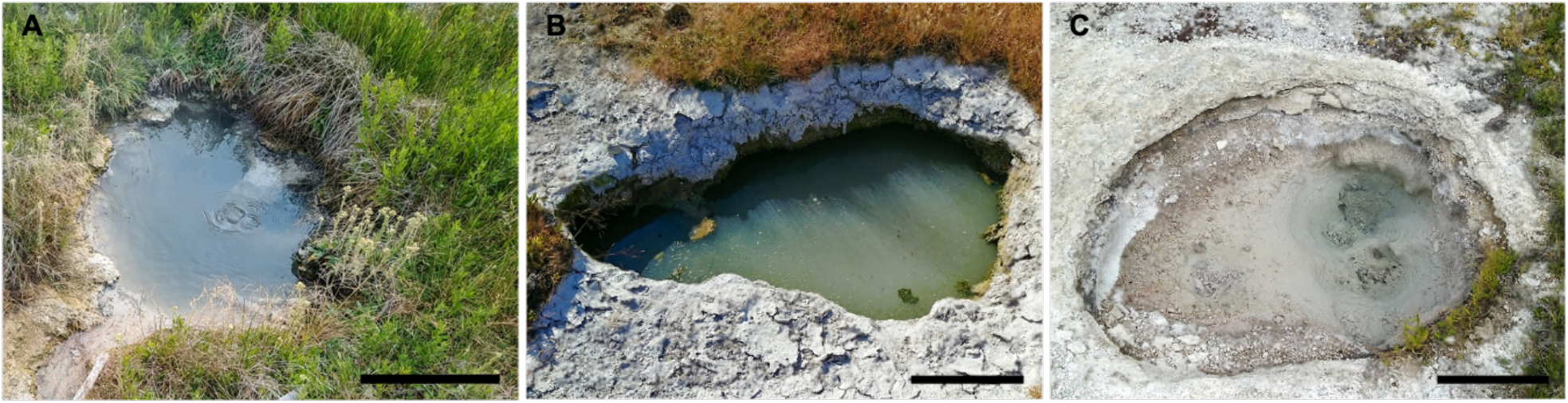
Photographs of main study sites. (A) LCB003, 2019-07-24, (B) LCB019, 2019-07-25, and (C) LCB024, 2019-07-23. Scale bar indicates 30 cm.

**SI Figure 5.**
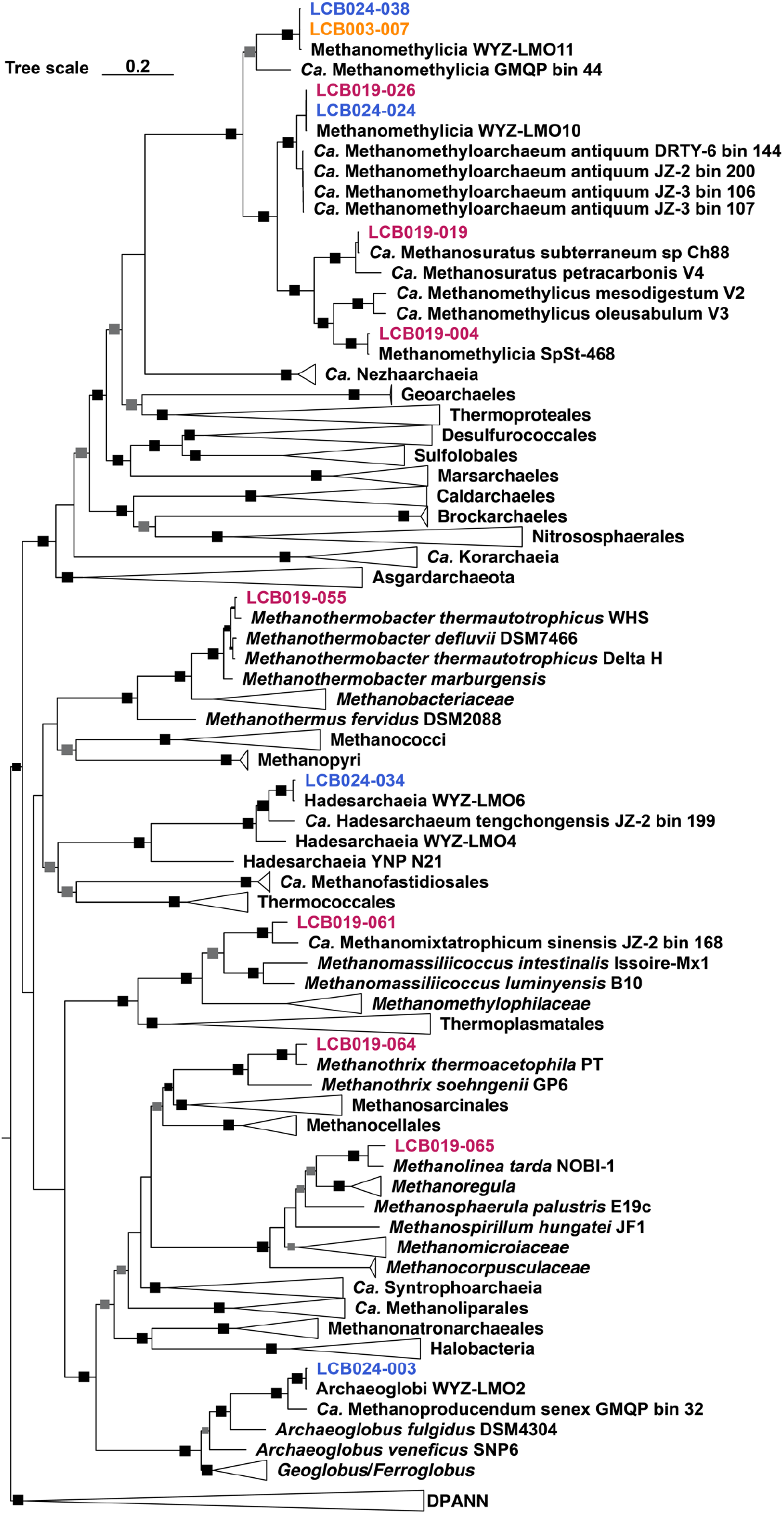
Phylogenetic affiliation of Mcr-encoding MAGs. Expanded maximum-likelihood tree, inferred with IQtree and the best-fit LG+C60+F+G model, from the concatenated alignment of 18 conserved arCOGs (SI Table 4). Squares indicate ultrafast bootstrap values of 100% (black) and 95-99% (gray).

**SI Figure 6.**
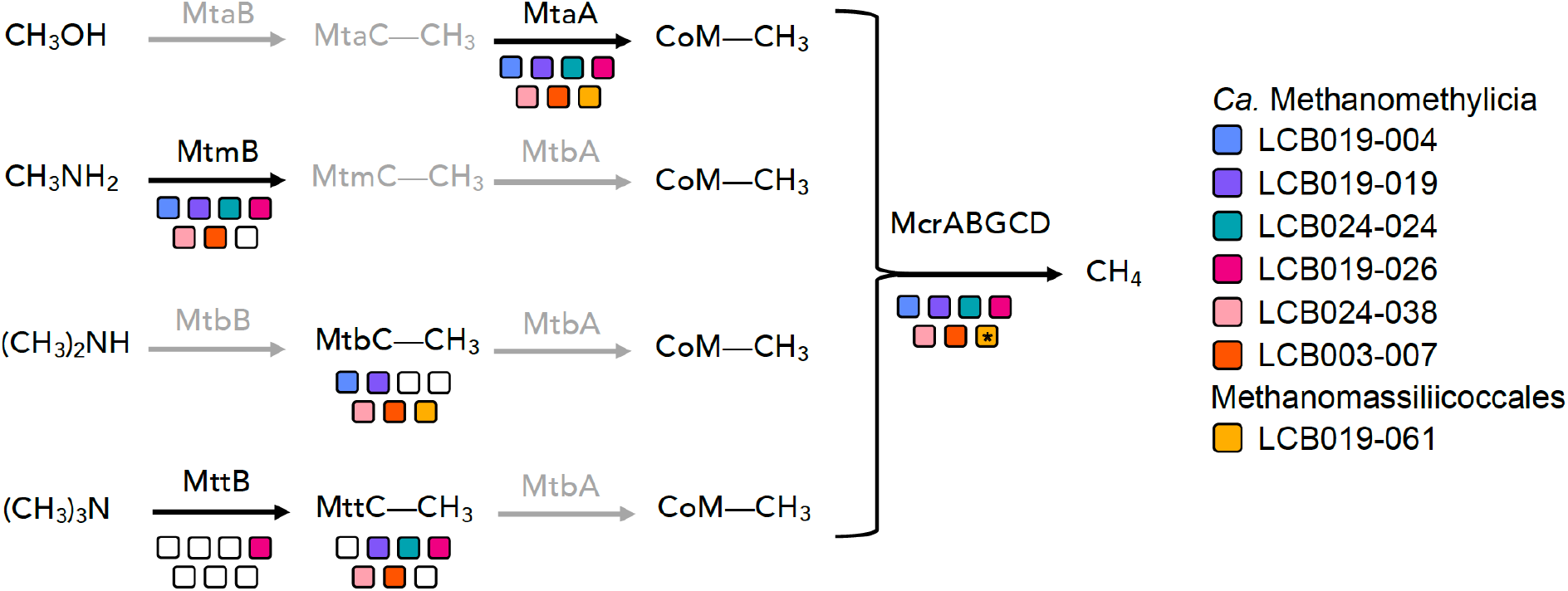
Proposed pathways for methylotrophic methanogenesis in MAGs from this study. Colors indicate individual MAGs affiliated with *Ca*. Methanomethylicia and Methanomassiliicoccales. Filled squares indicate methyltransferases and corrinoid proteins detected in the MAG. Shaded in gray are methyltransferases and corrinoid proteins not detected in any MAG. * denotes absence of McrG in LCB019-061. See SI Data 3 for details on protein abbreviations.

**SI Figure 7.**
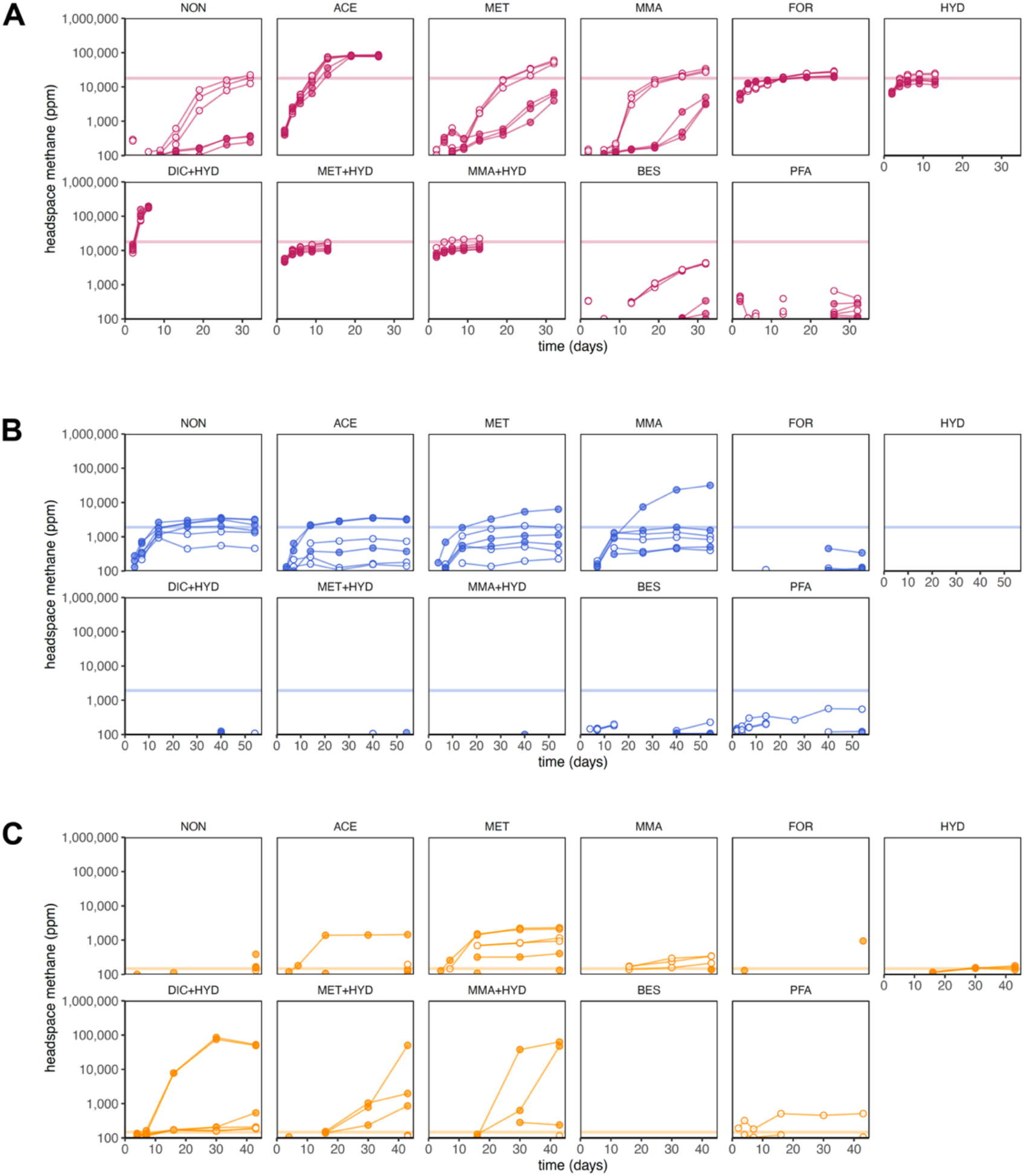
Development of methane concentrations in the headspace of mesocosms. (A) LCB019, (B) LCB024 and (C) LCB003. Open circles, without antibiotics; filled circles, with antibiotics. Horizontal line: average maximum methane produced in controls without substrate amendment and without antibiotics. Detection limit: 100 ppm. Abbreviations: NON, no substrate amendment; FOR, formate; ACE, acetate; MET, methanol; MMA, monomethylamine; HYD, hydrogen; MMA+HYD, monomethylamine plus hydrogen; MET+HYD, methanol plus hydrogen; DIC+HYD, dissolved inorganic carbon (HCO_3_^-^+CO_2_) plus hydrogen; BES, bromoethanesulfonate (methanogenesis inhibitor); PFA, paraformaldehyde (killed control).

**SI Figure 8.**
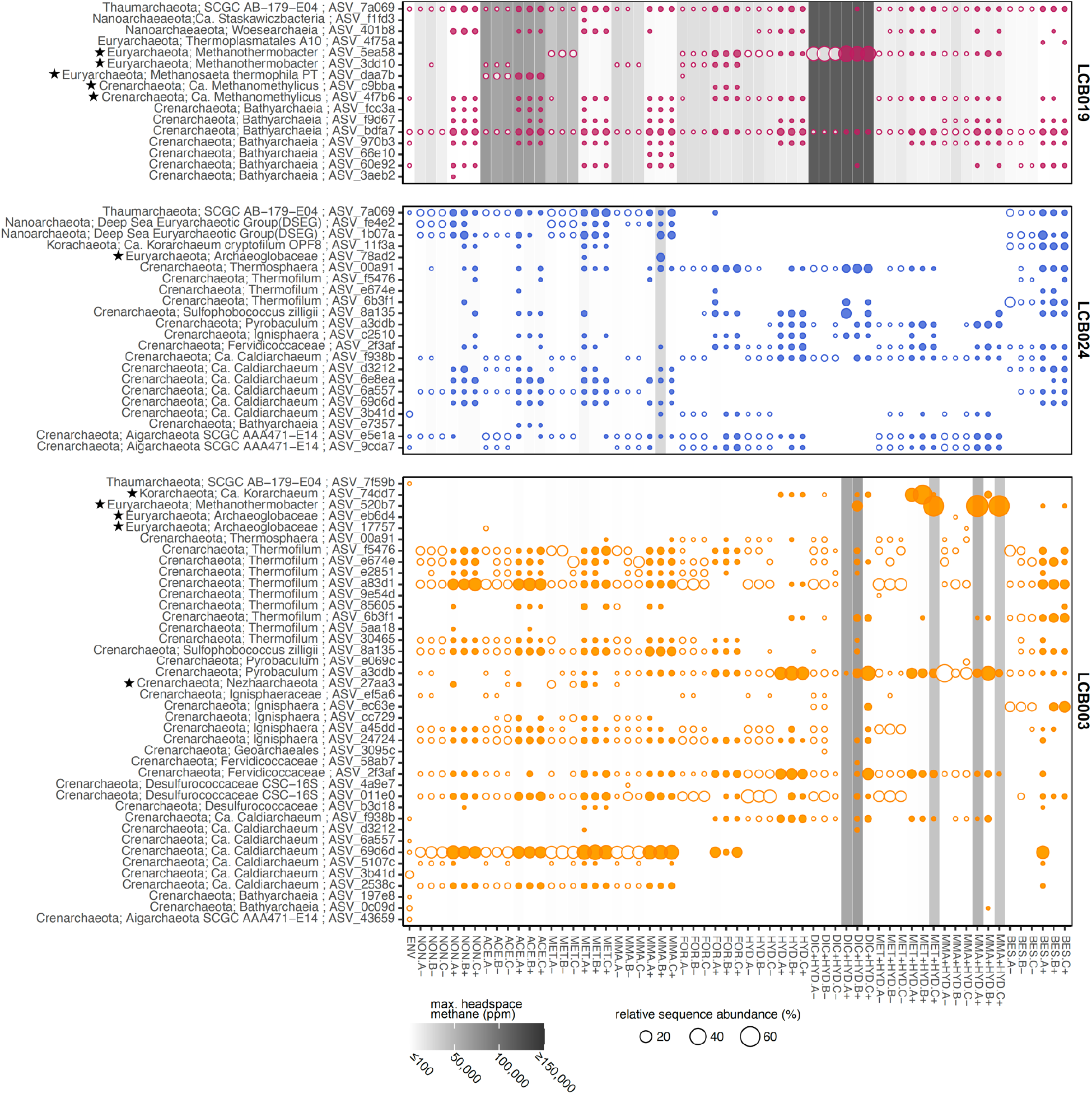
Relative sequence abundance of archaeal ASVs detected at >1% in mesocosms. Stars indicate ASVs identified as affiliated with Mcr-encoding archaea discussed in this study. Open circles, without antibiotics; filled circles, with antibiotics. Color coding by site: orange, LCB003, magenta, LCB019, blue, LCB024. Circle size proportional to relative sequence abundance. Gray shading indicates maximum methane concentration detected in mesocosm headspace. Abbreviations: ENV, environmental sample; NON, no substrate amendment; FOR, formate; ACE, acetate; MET, methanol; MMA, monomethylamine; HYD, hydrogen; MMA+HYD, monomethylamine plus hydrogen; MET+HYD, methanol plus hydrogen; DIC+HYD, dissolved inorganic carbon (HCO_3_^-^+CO_2_) plus hydrogen; BES, bromoethanesulfonate (methanogenesis inhibitor); PFA, paraformaldehyde (killed control).

**SI Figure 9.**
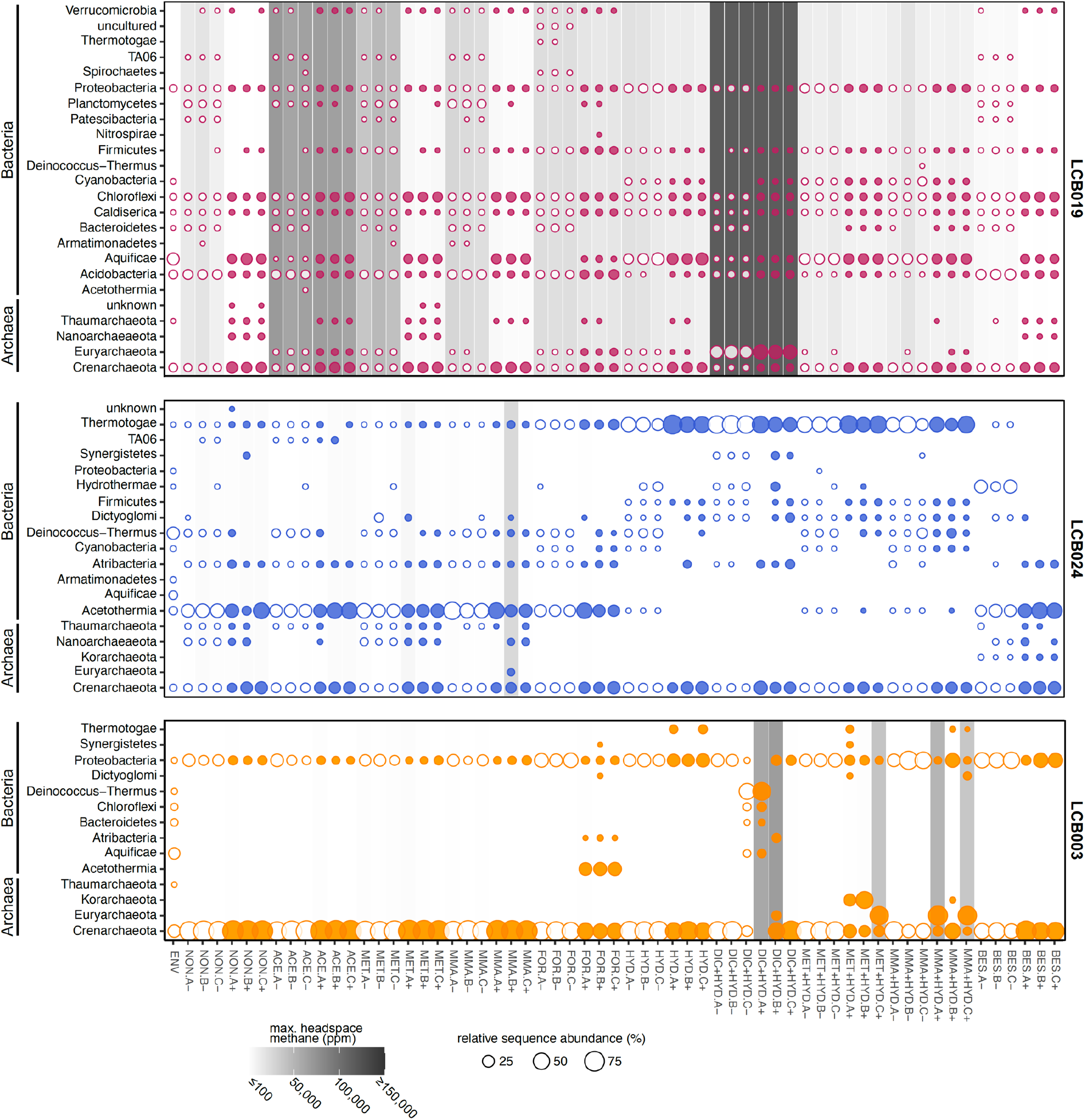
Relative sequence abundance of phyla detected at >3% in mesocosms. Open circles, without antibiotics; filled circles, with antibiotics. Color coding by site: orange, LCB003, magenta, LCB019, blue, LCB024. Circle size proportional to relative sequence abundance. Gray shading indicates maximum methane concentration detected in mesocosm headspace. Abbreviations: ENV, environmental sample; NON, no substrate amendment; FOR, formate; ACE, acetate; MET, methanol; MMA, monomethylamine; HYD, hydrogen; MMA+HYD, monomethylamine plus hydrogen; MET+HYD, methanol plus hydrogen; DIC+HYD, dissolved inorganic carbon (HCO_3_^-^ +CO_2_) plus hydrogen; BES, bromoethanesulfonate (methanogenesis inhibitor); PFA, paraformaldehyde (killed control).

## Extended Materials and Methods

### Physicochemical measurements, aqueous geochemistry, and elemental analysis

Prior to removing any sediment or microbial mat material from geothermal features, pH and temperature were recorded using a thermocouple (Fluke 52 II; Everett, WA) and a portable pH meter (ThermoFisher Orion Star A329 with electrode 8107UWMMD, Accumet AP63 with electrode AP50A, Accumet AP71 with electrode AP55, Waltham, MA), and geothermal water was collected for analysis of aqueous geochemistry and elements. Water samples were filtered through a pre-rinsed 0.22 µm PES syringe filter directly into sterile screw-cap plastic tubes and carbon-free glass vials sealed headspace-free. Blanks were prepared with ultrapure water (Barnstead, ThermoFisher, 18.2 MΩ). Samples and blanks were stored at 4 °C until analysis by the Environmental Analytical Lab (Montana State University). An aliquot was acidified with tracemetal grade HNO_3_ (20%) to a final concentration of 5% and analyzed using inductively coupled plasma optical emission spectroscopy (ICP-OES) (SpectroBLUE, Kleve, Germany) for total dissolved Ag, Al, As, Ba, Be, Bi, Ca, Cd, Co, Cr, Cu, Fe, Mg, Mn, Mo, Ni, Pb, Sb, Se, Sr, Ti, Tl, V, Zn, B, K, Li, Na, S, Si. An unacidified aliquot was analyzed for SO_4_^2-^, Cl^-^, and F^-^ using a Dionex (Sunnyvale, CA, USA) ICS-2100 ion chromatograph with AS18 column and NO_2_^-^, PO_4_^3-^, and NH_4_^+^ using a segmented flow analyzer (Seal Analytical, Southampton, UK). Samples collected in carbon-free glass vials were analyzed for total carbon (TC), inorganic carbon (IC), non-purgeable organic carbon (NPOC), and total nitrogen (TN) using a carbon analyzer (Shimadzu, Kyoto, Japan).

Dissolved oxygen (DO) was determined using a closed-headspace modification of the Winkler protocol (1). Briefly, 60 mL of geothermal water were collected with a 60 mL syringe that was immediately capped with a rubber septum to prevent gas exchange with the atmosphere. Needles and syringes were used to inject 0.4 mL of 2.15 M MnSO_4_ and 0.4 mL alkali-iodide-azide solution (12 M NaOH, 0.869 M KI, 0.15 M NaN_3_). The sample was thoroughly mixed by inversion and the formed floc was allowed to settle for 3-5 minutes. This mixing process was repeated twice before 0.4 mL concentrated (18 M) H_2_SO_4_ were added, and the floc completely dissolved by mixing. 30 mL of this mixture were titrated to colorlessness with 10.1 mM sodium thiosulfate to determine DO.

Total dissolved sulfide (H_2_S(aq)) was measured using the amine-sulfuric acid method (2). For this, 7.5 mL of geothermal water were collected with a 10 mL syringe and added into triplicate glass tubes pre-filled with 0.5 mL of amine sulfuric acid reagent (Ricca Chemical). Approximately 0.15 mL (4 drops) of 15 M FeCl_3_ were immediately added and the sample was mixed by slowly inverting the test tube once. After 3-5 minutes, 1.6 mL of 3.8 M (NH_4_)_2_PO_4_ were added and the sample was stored in the dark until absorbance was measured at 664 nm using a BioMate 3S spectrophotometer or Ocean Optics USB2000 spectrophotometer within 24 hours of sample collection.

The ferrozine assay was used to measure total and ferrous iron (Fe, Fe(II)) (3). Briefly, 5 mL of geothermal water were filtered through a 0.22 µm pore size PES syringe filter into three replicate tubes. 250 µL of a 10% w/v hydroxylamine solution were added for the Fe(II) assay before 250 µL of 4.9 mM ferrozine reagent and 625 µL ammonium-acetate buffer (3) were added for both Fe total and Fe(II) assays. After dilution to 12.5 mL with distilled water, samples were stored in the dark until absorbance was measured at 562 nm using a BioMate 3S spectrophotometer or Ocean Optics USB2000 spectrophotometer within 24 hours of sample collection.

To determine dissolved methane (CH_4_(aq)), geothermal water was collected using a peristaltic pump with an in-line 142 mm diameter, 0.2 µm pore size hydrophilic polycarbonate membrane filter (Millipore). 160 mL serum bottles were purged with three pour volumes of geothermal water, capped without headspace using butyl rubber stoppers, secured with aluminum rings, and stored upside-down at 4 °C. Prior to the analysis of dissolved gases, samples were weighed before and after 30 mL of liquid was removed, headspace replaced with ambient air, and incubated on a shaker at 60 rpm for two hours at room temperature. From the headspace, 1 mL was withdrawn using a gas tight syringe and injected into an SRI Multiple Gas Analyzer #5 gas chromatograph equipped with a Haysep D column (6 foot), a thermal conductivity detector, and a flame ionization detector with a methanizer (carrier gas: N_2_ at 13 psi, 50°C). Methane and mixed gas (CH_4_, CO_2_, CO, H_2_) standards (100 ppm, 10,000 ppm) were measured regularly. Concentrations of dissolved gases were calculated using Henry’s law and solubility equations (2, 4).

### *mcrA* and 16S rRNA gene amplification and amplicon sequencing

*mcrA* genes were amplified from DNA extracts of environmental samples and 16S rRNA genes were amplified from DNA extracts of mesocosm samples including the initial environmental material (environmental sample). DNA extracts from 500 µL sterile nuclease-free water were included as negative controls. DNA extracts were quantified using the Qubit high sensitivity assay (Invitrogen, Carlsbad, CA). Adapter sequences added to the gene-specific primers facilitated the preparation of Illumina amplicon libraries for sequencing as previously described (5).

The *mcrA* gene was amplified using the primer set mlas-mod-F/mcrA-rev-R (6). The PCR volume was 25 µL and consisted of 10 µL Invitrogen Platinum Taq II 2X Master Mix, 2 µL mlasmod-F primer (10 µM; final 0.8 µM), 2 µL mcrA-rev-R primer (10 µM; final 0.8 µM), 10 µL nuclease-free water, and 1 µL of template DNA (5 ng/µL). Thermocycler conditions for amplification were: 94°C for 4 min, followed by five touchdown cycles of 94°C for 30 sec, 60–1 °C for 45 sec, and 72°C for 30 sec, followed 30 cycles of 94 °C for 30 sec, 55 °C for 30 sec and 72 °C for 30 sec, and a final elongation step at 72 °C for 10 min (6).

Archaeal and bacterial 16S rRNA genes were amplified following the Earth Microbiome protocol with the updated primer set 515F and 806R (7-9). The PCR volume was 25 µL and consisted of 10 µL Invitrogen Platinum Taq II 2X Master Mix, 0.5 µL 515F primer (10 µM; final: 0.2 µM), 0.5 µL 806R primer (10 µM; final: 0.2 µM), 9 µL nuclease-free water, and 5 µL of template DNA (1 ng/µL). The thermocycler conditions were: 94 °C for 3 min followed by 28 cycles of 94 °C for 45 sec, 50 °C for 60 sec, and 72 °C for 90 sec before a final elongation step at 72 °C for 10 min.

A negative control with nuclease-free water as template was included in each PCR set. PCR products were checked for the expected length on a 1% agarose gel. *mcrA* and 16S rRNA gene amplicons were purified using AMPure XR beads (Beckman Coulter, Pasadena, CA) following the manufacturer’s protocol with a final elution volume of 40 µL nuclease-free water before a second PCR was performed to attach dual barcode indices and sequence adapters (Illumina). This PCR was performed in a 25 μL final volume with 5 μL purified amplicons 12.5 μL Invitrogen Platinum Taq II 2X Master Mix, 2.5 μL i5 primer (10µM; final: 0.25 μM), 2.5 μL i7 primer (10 µM; final: 0.25 μM), and 2.5 μL nuclease-free water. The PCR conditions were as follows: 95 °C for 3 min followed by 8 cycles of 95 °C for 30 sec, 55 °C for 30 sec, and 72 °C for 30 sec, followed by a final elongation step at 72 °C for 5 min. The amplicons were purified using AMPure XR beads and were quantified in triplicate reactions using the Quant-iT Picogreen dsDNA Assay (Invitrogen, Waltham, MA) and a Biotek Synergy H1 Hybrid microplate reader following manufacturers guidelines. Samples were pooled at 10 ng DNA each, concentrated using a QIAquick PCR purification spin column kit (Qiagen, Hilden, Germany) following the manufacturers protocol, and quantified with the Qubit high sensitivity assay (Invitrogen, Waltham, MA).

### Metagenome sequencing, read processing, assembly, and annotation

Truseq libraries were prepared at the JGI using low (10ng, LCB003.1) or regular (100ng, LCB019.1, LCB024.1) input quantities of DNA and were sequenced on the Illumina NovaSeq platform using the NovaSeq XP V1 reagent kits, S4 flow cell, following a 2 × 150 bp indexed run recipe. Raw metagenomic reads were processed according to the standard workflow employed by the JGI. Briefly, BBDuk (version 38.08; https://bbtool.jgi.doe.gov) was used to remove contaminants, trim reads that contained adapter sequence, right quality trim reads where quality drops to zero and remove reads that contained four or more ‘N’ bases, had an average quality score across the read less than 3 or had a minimum length <= 51 bp or 33% of the full read length. Reads mapped with BBMap (v37.78) to masked human, cat, dog, and mouse references at 93% identity were removed as were reads aligned to common microbial contaminants (JGI SOP 1077). Trimmed and screened paired-end reads were error corrected with bfc (v. r181) with options: -1, - s 10g, -k 21, t 10 (Li, 2015) and then assembled using SPAdes 3.11.1 (10) with options -m 2000, -k 33,55,77,99,127 -meta. Coverage information was generated using BBMap by mapping the entire filtered reads dataset against the final assembly. Annotation of the final assembly was performed with JGI’s Integrated Microbial Genomes & Microbiomes Expert Review (IMG/M-ER) pipeline v7 (11). Assembled scaffolds ≥2,000 bp were binned with six implementations of four different programs including, Maxbin v2.2.4 (12), Concoct v1.0.0 (13), Metabat v2.12.1 (with and without coverage) (14), and Autometa v1 (with bacterial and archaeal modes, and Machine Learning step, but omitting the taxonomy assignment step) (15). Bins generated from each method were dereplicated, aggregated, and scored with DAS_Tool (16).

## Extended Discussion

### Energy conservation mechanisms of Mcr-encoding archaea

Both LCB019-065 (*Methanolinea*) and LCB019-055 (*Methanothermobacter*) encode heterodisulfide reductase (HdrABC) and either F_420_-non-reducing hydrogenase (MvhADG; LCB019-055) or F_420_-reducing hydrogenase (FrhAG) and MvhD (LCB019-065) allowing for two distinct mechanisms of heterodisulfide (CoM-S-S-CoB) reduction. In LCB019-055, heterodisulfide reduction could be mediated by an electron bifurcating HdrABC/MvhADG complex as is common for other hydrogenotrophic methanogens (17). Conversely, in LCB019-065, HdrABC could form an electron bifurcating complex with FrhAG and MvhD to reduce heterodisulfide as described in some members of the Methanomicrobiales (17-19).

LCB019-064 (*Methanothrix*) encodes HdrDE and a Fpo-like complex (*fpoABCDHIJKLMN*, lacking *fpoFO* for binding and oxidizing F_420_), which would allow for heterodisulfide reduction by HdrDE with electrons supplied by the Fpo-like complex. The [NiFe] hydrogenase Frh, energy-converting hydrogenase (Ech), Na^+^-translocating ferredoxin:NAD^+^ oxidoreductase (Rnf), and methanophenazine-reducing hydrogenase (Vho) were not encoded. This suggests that energy conservation in LCB019-064 may proceed as proposed for *Methanothrix thermophila* (20) and differs from the energy conservation mechanisms of other aceticlastic methanogens within the genus *Methanosarcina*.

Methanomassiliicoccales MAG LCB019-061 encodes HdrABC, MvhADG, and a partial Fpo-like complex (20), which is consistent with the energy conservation complexes found in H_2_-dependent methylotrophic methanogens (21). An electron bifurcating HdrABC/MvhADG complex (22, 23), could couple the oxidation of H_2_ with the reduction of heterodisulfide and ferredoxin. Interestingly, multiple copies of the *hdrA* (x14), *hdrB* (x4), and *hdrC* (x4) genes were identified in LCB019-061 while *hdrD* was not detected (SI Data 3). Although other Methanomassiliicoccales MAGs encode HdrD, the Fpo-like complex could function without an association to HdrD as in *Methanothrix* sp. (20, 24). The oxidation of reduced ferredoxin by the membrane bound Fpo-like complex would generate a proton gradient for ATP synthesis via a V/A-type ATP synthase.

The energy conservation complexes encoded in the six *Ca*. Methanomethylicia MAGs (LCB003-007, LCB019-004, LCB019-019, LCB019-026, LCB024-024, and LCB024-038) were similar to those in related MAGs (25). Genes encoding complexes involved in establishing a proton gradient, including the Fpo-like complex (20) and an energy-converting hydrogenase B (Ehb) complex, were detected in all six MAGs (26). The generated proton gradient would drive ATP synthesis via a V/A-type ATP synthase, encoded in all MAGs. *hdrABC* and *mvhADG* were absent in the MAGs suggesting coenzymes M and B are regenerated in an alternative way. Previously proposed mechanisms of heterodisulfide reduction encoded in the six *Ca*. Methanomethylicia MAGs include reduction via an interaction between HdrD and the Fpo-like complex (24), a Fpo-HdrD-GlcD complex (25), or energy-converting complex D (Ehd) formed by a membrane-bound [NiFe] hydrogenase and HdrBC (27). All six MAGs contained 1-3 copies of *hdrD* and *glcD*, however the Ehd complex was only encoded in four MAGs (LCB003-007, LCB019-026, LCB024-038, and LCB019-019 (partially)).

LCB024-003 (Archaeoglobi) encodes two mechanisms for coenzymes M and B regeneration. LCB024-003 contains *mvhADG, hdrD*, and three copies of *hdrA* (one upstream of *mvhADG). hdrBC* were not detected; however, HdrD has been proposed to substitute for HdrBC in heterodisulfide reduction in other Mcr-encoding Archaeoglobi MAGs (28, 29). LCB024-003 also encodes a fused HdrDE and a membrane-bound F_420_H_2_:quinone oxidoreductase (Fqo) complex. In Archaeoglobi isolates, the Fqo complex has been shown to reduce menaquinone instead of methanophenazine (30), which is characteristic of methanogens containing an Fpo complex (31). HdrDE contains two cysteine-rich domains (CCG) and is homologous to the membrane bound complex DsrMK, which is inferred to reduce a cytoplasmic cysteine disulfide (Cys-S-S-Cys) in DsrC during dissimilatory sulfate reduction (32). Similarly, HdrDE could couple the periplasmic oxidation of menaquinol, generated from Fqo, to the cytoplasmic reduction of the disulfide bond in CoM-S-S-CoB. A replacement of HdrDE for DsrMK has been shown in *Archaeoglobus fulgidus* VC16 (33) and has been proposed for other Mcr-encoding Archaeoglobi MAGs (29).

